# An epigenetic bifunctional that toggles between transactivation and repression

**DOI:** 10.64898/2026.03.17.712509

**Authors:** Ananthan Sadagopan, Maximilian Carson, Eriks J. Zamurs, Shanivi Srikonda, Cary N. Weiss, Michael J. Bond, Anurag Sodhi, Katherine A. Donovan, Julia K. Ryan, Eric S. Fischer, Kimberly Stegmaier, Srinivas R. Viswanathan, Benjamin L. Ebert, William J. Gibson

## Abstract

The targeted modulation of gene expression with bifunctional small molecules enables the precise control of cellular and biological processes. To screen for ligands that could be used to induce gene expression, we conjugated the high affinity FKBP(F36V) binder, AP1867, to known high-affinity binders of activating epigenetic machinery. We tested these bifunctionals in a FKBP(F36V)-tagged transcription factor reporter system and found bifunctional induced transactivation is relatively common, being observed for bifunctionals with BET ligand JQ1, p300/CBP ligand GNE-781, CDK9 ligand SNS-032, and BRD9 ligand iBRD9. aTAG-2 (mAP1867-C8-GNE781) was identified as the strongest and most potent transactivator, possessing single-digit nanomolar activity. When tested in models where oncogenic RNA binding protein-transcription factor fusion proteins have been FKBP(F36V)-tagged, we unexpectedly observed rapid collapse of the fusion transcriptional program. In a tagged Ewing sarcoma model, aTAG-2 exhibits at least three distinct mechanisms of action: i) RIPTAC mediated p300/CBP inhibition, ii) ubiquitination- and ternary complex-dependent EWS/FLI degradation, and iii) replacement of p300 with CBP at EWS/FLI bound chromatin loci. Together, these data establish bifunctionals targeting p300/CBP that toggle between a program of ultra-potent transactivation and repression depending on cellular context. Overall demonstrating that induced proximity with a given ligand does not encode a fixed functional outcome.

## Introduction

Bifunctional small molecules can reprogram biology by inducing proximity between proteins, enabling event-driven mechanisms that go beyond classical occupancy-based inhibition. Early chemical inducer of dimerization (CID) systems established that small molecules can control protein interactions in cells,^1^ and this logic has since expanded into multiple proximity-enabled modalities, including targeted degradation (PROTACs),^2–4^ molecular glue-like neosubstrate recruitment,^5–9^ and a variety of enzymatic and non-enzymatic events.^10–18^ Together, these advances highlight proximity as a general chemical strategy to redirect endogenous cellular machinery toward new outcomes.

For modulating transcription, the concept that bringing the right protein to the right location can be sufficient to regulate gene expression has long-standing roots in the study of synthetic transcription factors. Prior work has often shared a modular theme in which a DNA binding component, such as a minor-groove binding polyamide/triplex-forming oligonucleotide, is linked to a minimal transcriptional activation module.^19–27^ Other work has demonstrated hyper-activation of existing small molecule ligands for DNA binding proteins for example by tethering a transactivating moiety to dexamethasone.^28,29^ In parallel, small-molecule transcriptional activation domain mimics demonstrated that compact synthetic ligands can replace or tune activating functions and thereby expand the chemical space of transcriptional control.^30,31^ More recently, work in this space has been accelerated by modular tagging systems and the development of high-affinity ligands for endogenous coactivator machinery. The FKBP^F36V^ tag and its ultra-high affinity ligands (AP1867 derivatives) enable robust, selective engagement of engineered targets^32^ and have been broadly deployed for rapid target depletion through the dTAG platform.^33,34^

In the gene-regulation setting, several modern strategies demonstrate that induced proximity can drive transcription when the recruited effector is appropriately chosen. One class of approaches uses programmable DNA targeting to localize small-molecule recruitment systems to defined genomic loci; for example, dCas9-coupled recruitment platforms enable dose-dependent gene activation through chemically controlled effector engagement.^35^ A second strategy uses synthetic transcription elongation factors (SynTEFs) that engage BRD4 to promote productive transcription through otherwise repressed chromatin environments.^25^ Related bifunctional designs directly tether transcriptional coactivators to DNA-bound regulators. Transcriptional chemical inducers of proximity (TCIPs) recruit factors such as BRD4, CDK9, or p300/CBP to the oncogenic repressor BCL6, thereby rewiring transcriptional programs and inducing context-dependent oncogenic phenotypes.^36–38^ Similar principles have been extended to fusion-oncogene contexts using epigenetic bifunctionals that recruit EWS/FLI to re-activate repressed BCL6 targets.^39^ More broadly, induced proximity can also be tuned to generate selective cellular vulnerabilities. RIPTAC frameworks, for example, exploit differential target abundance to create cell-state-dependent therapeutic windows for proximity-based mechanisms.^18,40^

Despite these advances, it remains difficult to compare the relative activating strength of different recruited transcriptional or epigenetic machineries within a common experimental framework, and to predict when forced proximity will produce productive activation versus disruption of an already-engaged transcriptional program, particularly in oncogenic fusion settings where transcription factor levels and coactivator balance are tightly constrained. To address these challenges, we systematically explored bifunctional “activating TAG” (aTAG) molecules that couple FKBP12^F36V^ engagement to diverse endogenous activating machineries, first in a defined IRF1 reporter system and then in FKBP^F36V^-tagged oncogenic fusion models. This strategy allowed us to examine how effector identity and cellular context shape proximity-driven transcriptional outcomes, and to test whether proximity-mediated hyperactivation or coactivator rewiring can perturb oncogene-addicted states.

## Results

### Small molecule recruitment of activating epigenetic machinery to a FKBP^F36V^-tagged transcription factor induces gene activation

To create a system to assess bifunctional small molecule-mediated transactivation, we integrated an interferon regulatory factor 1 (IRF1)-responsive luciferase reporter and lentiviral doxycycline-inducible interferon regulatory factor 1 (IRF1)-FKBP^F36V^-mCherry fusion construct into U2OS cells. IRF1 is a representative human transcription factor, with a defined consensus sequence, well-characterized transcriptional signature, constitutive nuclear localization, transactivation domain, and can bind DNA as both a monomer or dimer. Fusion of FKBP^F36V^ enabled us to ligand IRF1 with the ultra-high affinity FKBP^F36V^ binder *meta*-AP1867 (K_d_ ∼ 94 pM).^32^

We next synthesized a series of bifunctional small molecules binding to FKBP^F36V^ tag and endogenous activating machinery (**Fig. 1A**). We amidated AP1867 with a short linker (C8, PEG2) and coupled the resulting molecules with binders of activating epigenetic machinery, which we resynthesized with solvent-exposed amines/acids/alkynes; we preferred binders with defined exit vectors in PROTAC literature (**Table 1**).

**Table 1.**
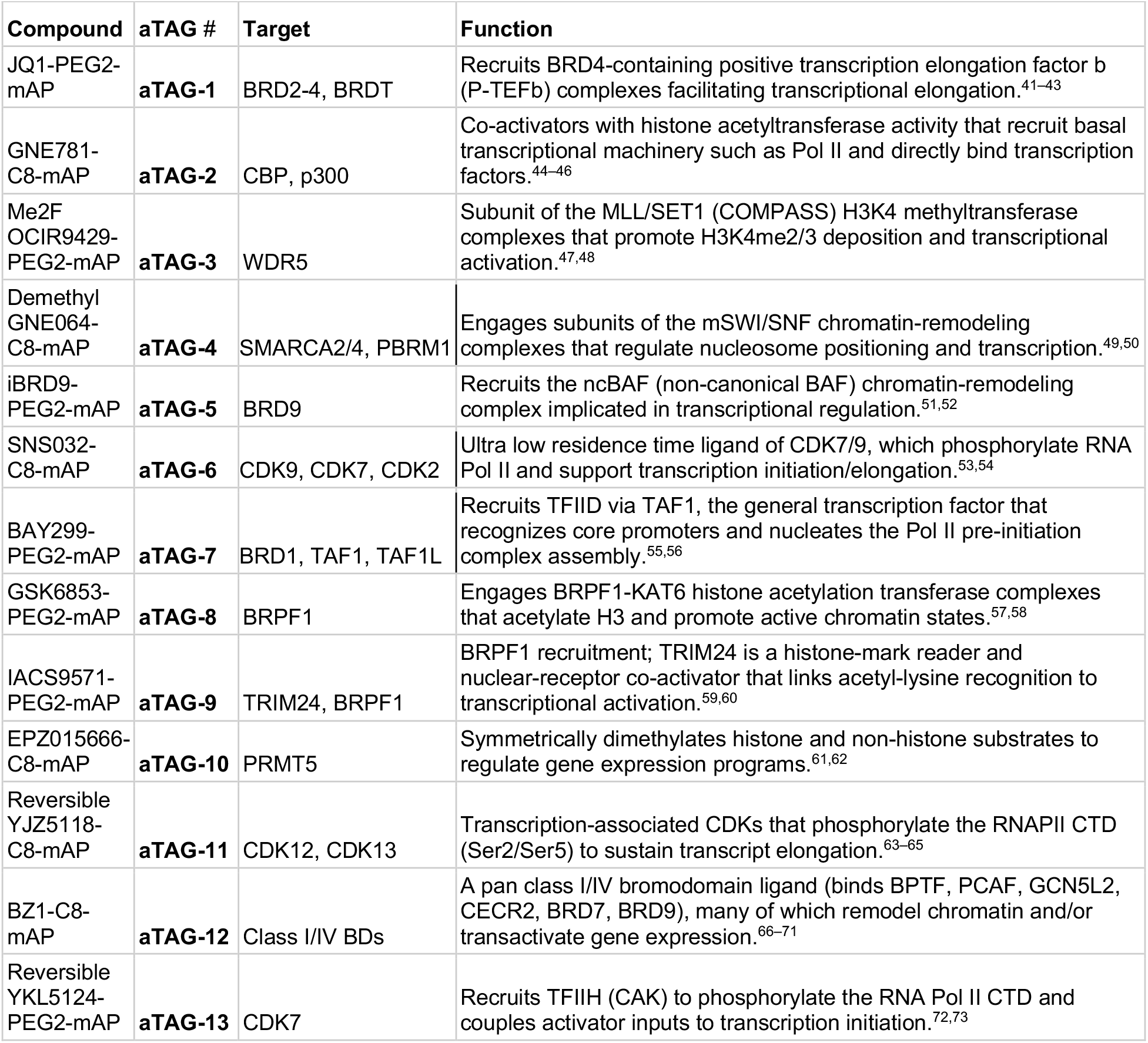
Library of aTAG molecules synthesized.

**Fig. 1.**
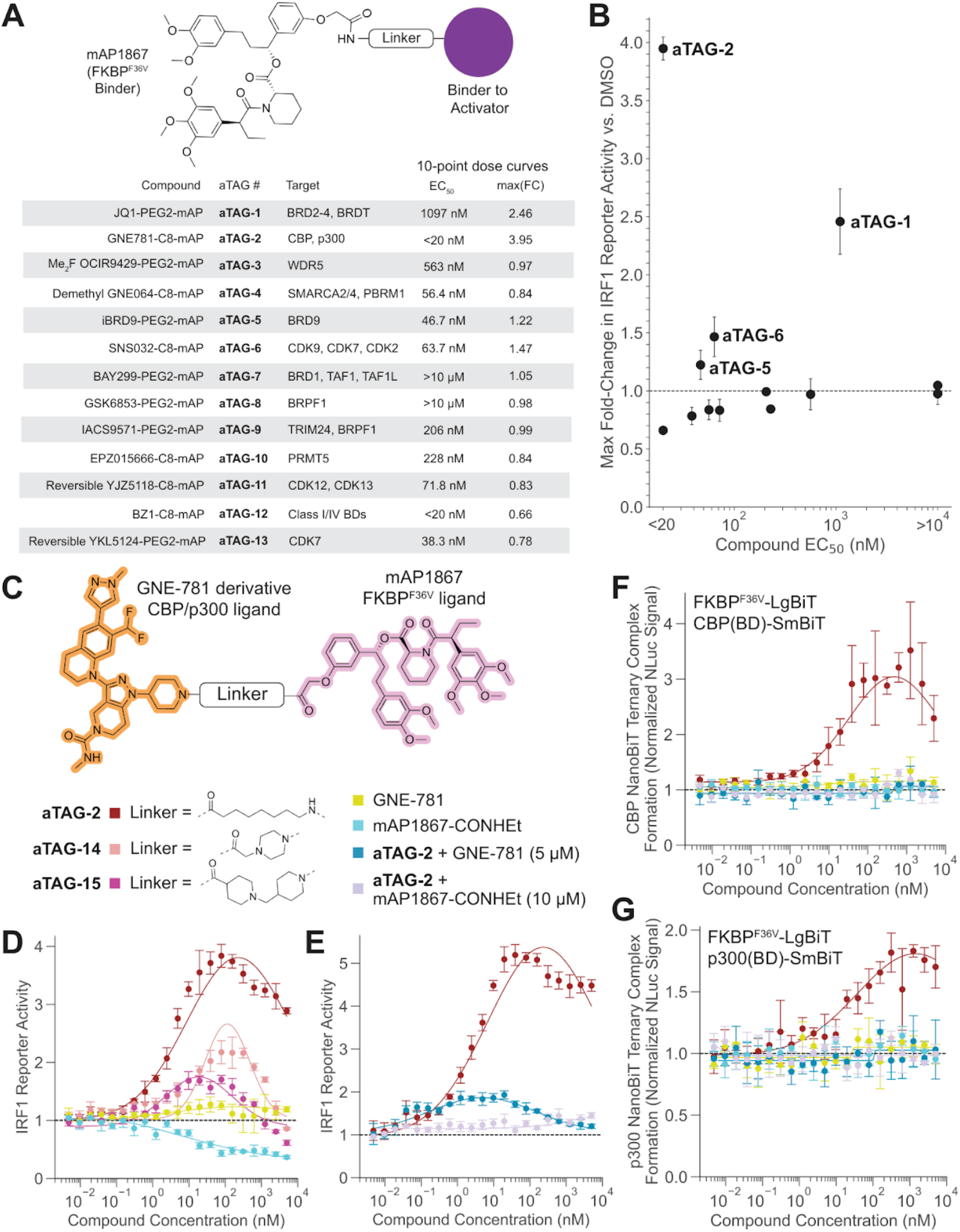
Screening of epigenetic bifunctionals yields CBP/p300 ligand GNE-781 as the strongest transactivator. A) Chemical architecture of a representative aTAG showing the FKBP^F36V^ ligand (AP1867), the activator ligand (purple) and a linker; lower table summarizes the 13 aTAG molecules synthesized, the chromatin effector they recruit, reporter EC_50_ values (note: for biphasic curves such as aTAG-5/aTAG-6, this represents the interpolated dose with maximal activation) and maximal activation (fold-change vs. untreated at a single dose) from 10-point dose titration in an IRF1 reporter system (48h treatment); U2OS reporter system stably expressing 1) a plasmid containing 4 repeats of the IRF1 consensus sequence upstream of firefly luciferase, 2) a plasmid containing doxycycline-inducible IRF1-FKBP^F36V^-mCherry in pLIX403. B) Scatter-plot of maximal activation versus potency for all compounds after 48h treatment. C) Chemical structures of linker series tested for GNE-781 bifunctionals. D) Extended dose curves of AP1867-GNE781 bifunctionals or binders alone (GNE-781, mAP1867-CONHEt) in IRF1 reporter system (normalized FLuc luminescence at 48 hours). E) Extended dose curves of aTAG-2 with binder competition in IRF1 reporter system (normalized FLuc luminescence at 48 hours). F-G) Normalized NLuc luminescence from NanoBiT experiment in 293T cells, titrated with aTAG-2 ± competitor or binder alone for 24 hours. NanoBiT constructs used were FKBP^F36V^-LgBiT and SmBiT fused to the C-terminus of either the CBP (F) or p300 (G) bromodomain.

The majority of bifunctional molecules induced no gene activation in the reporter (9/13, 69.2%) with four molecules displaying transactivation capability: **aTAG-1** (JQ1, BET BDs), **aTAG-2** (GNE-781, p300/CBP), **aTAG-5** (iBRD9, BRD9), and **aTAG-6** (SNS-032, CDK7/9). Apart from iBRD9, these ligands have demonstrated transactivation in other contexts^25,35–38^; a BRD9 molecular glue was also recently shown to be transactivating.^74^ **aTAG-5** and **aTAG-6** were weak inducers of gene expression, with **aTAG-1** being considerably stronger, and **aTAG-2** being both the most efficacious and potent at 48 hours (**Fig. 1B, Fig. S1A**).

### aTAG-2 displays behavior characteristic of a bifunctional small molecule

Encouraged by our initial data with **aTAG-2**, we attempted to rigidify the C8 alkyl linker (creating **aTAG-14** and **aTAG-15**). These changes failed to increase potency or E_max_ in our reporter (**aTAG-2**: EC_50_ ∼ 1-5 nM, E_max_ ∼ 4-5 fold, we did not perform further linker optimization given the high baseline potency of **aTAG-2**). Hook effects were observed for all molecules in the linker series, being least pronounced for **aTAG-2**. The binders alone, GNE-781 or AP1867 ethyl amide (mAP1867-CONHEt) did not activate the reporter **(Fig. 1C-D**). The transactivation capability of **aTAG-2** was attenuated by excess GNE-781 (5 μM) or AP1867 ethyl amide (10 μM) (**Fig. 1E**).

We next assessed ternary complex formation with the p300/CBP bromodomains and FKBP^F36V^ in a NanoBiT (split nanoluciferase) assay. We transfected constructs containing SmBiT-tagged bromodomains and LgBiT-tagged FKBP^F36V^ into 293T cells, and dose titrated molecules for 24 hours. We observed a dose-dependent reconstitution of nanoluciferase activity with the addition of **aTAG-2**, while binders alone (GNE-781, AP1867 ethyl amide) showed no increase in NanoBiT signal. **aTAG-2** was roughly equipotent for p300 and CBP ternary complex formation (p300 EC_50_ = 30 nM, CBP EC_50_ = 18 nM) and could be competed off with excess GNE-781 (5 μM) or AP1867 ethyl amide (10 μM) (**Fig. 1F-G**).

### aTAG-2 collapses the EWS/FLI transcriptional program in a Ewing sarcoma FKBP^F36V^ fusion model

Fusion oncogene expression is known to be tightly controlled in tumors driven by said alterations. Overexpression of a fusion or even knockout of an E3 ligase destabilizing a fusion can create significant fitness defects.^75–79^ Therapeutic overactivation (TOVER) is a recently contemplated therapeutic strategy for diverse oncogenic pathways, yet there remain few tools to study this, particularly for fusion oncogene driven cancers.^80,81^ We hypothesized that aTAG-2 could hyperactivate oncogenic fusions bearing the FKBP^F36V^ dTAG degron, thereby inducing a therapeutic overdose of oncogenic signaling. We focused on a Ewing sarcoma line (EWS502) where the endogenous fusion has been knocked out and FKBP^F36V^-HA-EWS/FLI was expressed at a physiological level; this cell line has been previously characterized, is responsive to dTAG-based degraders, and is dependent on an E3 ligase (TRIM8) destabilizing the fusion, thus making it an ideal model for such experiments.^34,39,77^

**aTAG-2** selectively inhibited proliferation of FKBP^F36V^-HA-EWS/FLI EWS502 cells while sparing the parental line at doses between 1-50 nM (IC_50_ = 2.33 nM in FKBP^F36V^-HA-EWS/FLI cells at 96 hours, **Fig. 2A**). Monovalent binders alone (GNE-781, AP1867 ethyl amide) did not substantially impact viability of FKBP^F36V^-HA-EWS/FLI EWS502 cells across this dose range, though high dose GNE-781 was toxic, consistent with reported p300/CBP dependency in this cell line.^82,83^ AP1867 ethyl amide (10 μM) was able to completely eliminate the toxicity induced by **aTAG-2** across this dose range, consistent with a bifunctional mechanism of action (**Fig. 2A**).

**Fig. 2.**
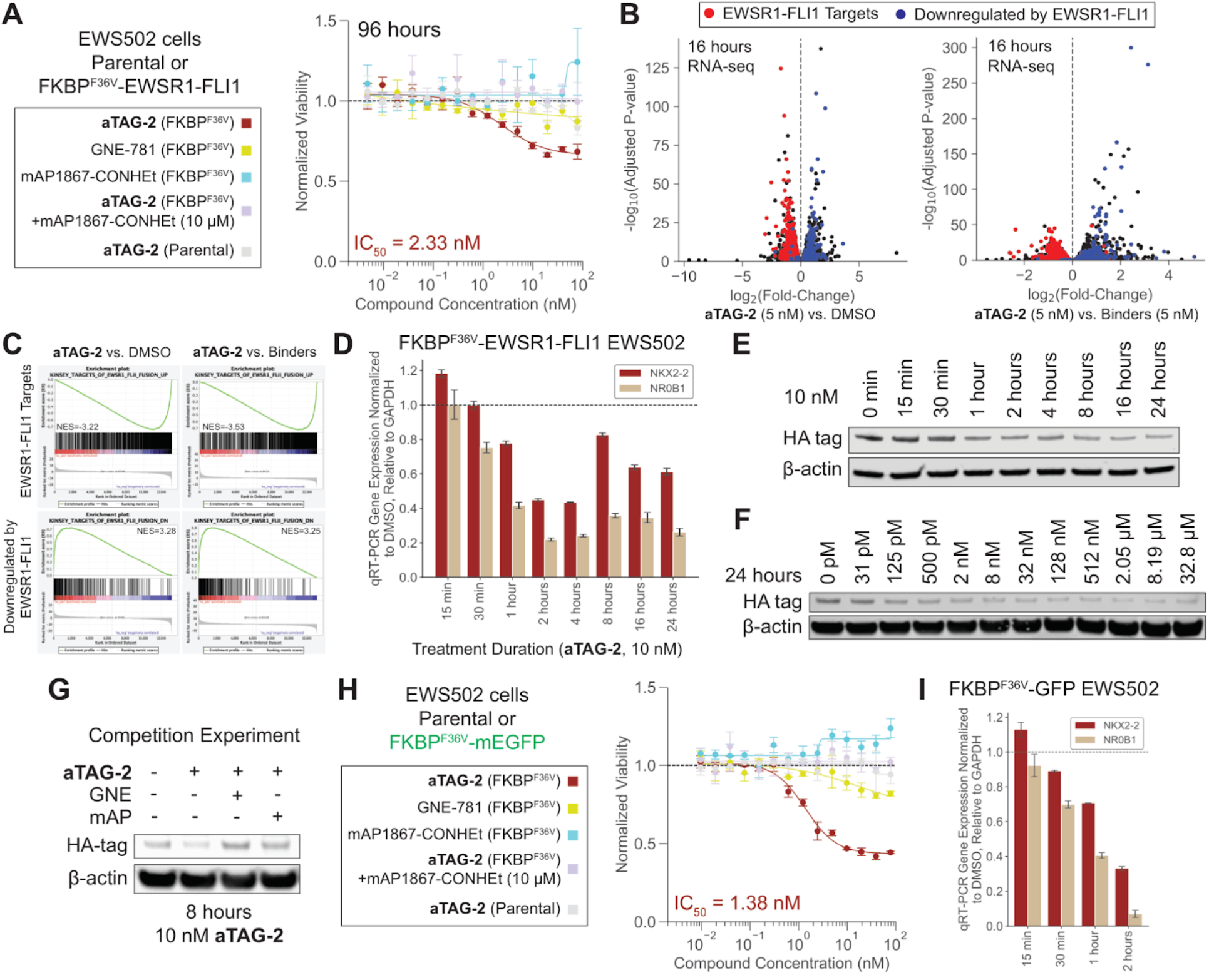
aTAG-2 is an ultra-potent ternary complex-dependent degrader and RIPTAC in a Ewing sarcoma with FKBP^F36V^-tagged EWS/FLI. A) Dose titration of aTAG-2 ± mAP1867-CONHEt or binders alone in FKBP^F36V^-HA-EWS/FLI or parental EWS502 cells. GNE-781 competition was not performed due to toxicity. Cell viability was read out at 96 hours by CTG. B) RNA-seq volcanoes (DESeq2) at 16 hours comparing aTAG-2 (5 nM) to either DMSO or binders (mAP1867-CONHEt + GNE-781 at 5 nM each). Kinsey et al. targets of EWSR1/FLI1 fusion UP are labelled in red (genes transactivated by EWS/FLI), DOWN labelled in blue (genes downregulated by EWS/FLI). C) Individual pre-ranked gene set enrichment plots across Kinsey et al. EWS/FLI gene sets from RNA-seq experiment in panel B. D) RT-qPCR for EWS/FLI targets (NKX2-2, NR0B1; normalized to GAPDH) vs. treatment duration of aTAG-2 (10 nM) in FKBP^F36V^-HA-EWS/FLI EWS502 cells. E-G) Western blot for HA-tagged EWS/FLI in FKBP^F36V^-HA-EWS/FLI EWS502 cells treated with aTAG-2 at 10 nM with time course (E), 24 hours with dose titration (F), or 8 hours at 10 nM with binder competition (mAP: 10 μM mAP1867-CONHEt, GNE: 5 μM GNE-781) (G). H) Dose titration of aTAG-2 ± mAP1867-CONHEt or binders alone in FKBP^F36V^-mEGFP or parental EWS502 cells. GNE-781 competition was not performed due to toxicity. Cell viability was read out at 96 hours by CTG. I) RT-qPCR for EWS/FLI targets (*NKX2-2, NR0B1*; normalized to *GAPDH*) vs. treatment duration of aTAG-2 (50 nM) in FKBP^F36V^-mEGFP EWS502 cells.

We performed RNA-seq following **aTAG-2** treatment (5 nM) for 16 hours and to our surprise, we observed downregulation of EWS/FLI targets and upregulation of genes normally repressed by EWS/FLI, using DMSO- or binder-treated (5 nM AP1867 ethyl amide + 5 nM GNE-781) cells as comparators. These changes were extremely robust, with literature reported EWS/FLI target/repressed gene sets^84^ consistently demonstrating absolute normalized enrichment scores > 3.2 (**Fig. 2B-C, Fig. S3A**). To understand the kinetics of transcriptional downregulation, we performed a RT-qPCR time course for EWS/FLI targets *NR0B1* and *NKX2-2*. Downregulation of both target genes occurred within 0.5-1 hour, reaching a maximal effect within 2 hours (**Fig. 2D**).

We also compared the changes in gene expression to previously generated RNA-seq following dual p300/CBP degradation (using a GNE781-based degrader, dCBP-1) in EWS502 cells.^82^ Unlike **aTAG-2**, dCBP-1 caused downregulation of both EWS/FLI targets (NES = -1.64) and genes normally repressed by EWS/FLI (NES = -1.91) (**Fig. S3B**). Together, these findings indicate that **aTAG-2** does not simply phenocopy global p300/CBP loss, but instead more selectively collapses the EWS/FLI transcriptional program.

We also profiled **aTAG-1, aTAG-5**, and **aTAG-6** in FKBP^F36V^-HA-EWS/FLI vs. parental EWS502 cells. **aTAG-5** did not impact cell viability at concentrations ≤ 10 μM. The remaining compounds selectively inhibited proliferation of FKBP^F36V^-HA-EWS/FLI EWS502 cells (**aTAG-1**: IC_50_ parental/FKBP^F36V^ = 1.34 μM/1.09 μM = 1.23x; **aTAG-6**: IC_50_ parental/FKBP^F36V^ = 2.29 μM/557 nM = 4.1x) albeit with smaller therapeutic windows and far less potently than **aTAG-2** (**Fig. S2A**). We profiled **aTAG-1** (250 nM) and **aTAG-6** (500 nM) by RNA-seq and observed evidence of significant EWS/FLI target gene repression for both compounds relative to DMSO-treated cells at 16 hours (**Fig. S2B-E**).

Together, these data show that while aTAG molecules are potent transcriptional activators in a transcriptional reporter system, they paradoxically decrease fusion oncoprotein transcriptional output in this Ewing sarcoma model.

### aTAG-2 acts as a degrader and RIPTAC in a Ewing sarcoma FKBP^F36V^ fusion model

Given the rapid decrease in EWS/FLI gene target expression, we hypothesized that **aTAG-2** may be inducing degradation of the fusion protein. Consistent, though incomplete, degradation of FKBP^F36V^-HA-EWS/FLI was observed within 1 hour in the presence of low doses of aTAG-2 (10 nM, **Fig. 2E**). Further dose titration revealed fusion degradation with doses as low as 125 pM after 24 hours (**Fig. 2F**). Compound-induced degradation could be competed off completely by GNE-781 (5 μM) or AP1867 ethyl amide (10 μM), with binders alone being inactive (**Fig. 2G, Fig. S3C**). Global proteomics analysis confirmed FLI1 degradation (note: wild-type, untagged EWSR1 is expressed in this cell line confounding readouts of EWSR1 degradation) (**Fig. S3D**).

We hypothesized that partial degradation alone was an incomplete explanation for the phenotypes we observed given dTAG-V1 induces complete degradation of the fusion in this cell line (as previously shown)^34^ with seemingly weaker effects on gene expression (**Fig. S3E**). We therefore considered the possibility that **aTAG-2** differentially inhibits CBP/p300 in FKBP^F36V^-tagged cells relative to parental cells, consistent with a RIPTAC-like mechanism. To test this, we generated an EWS502 cell line stably expressing FKBP^F36V^-mEGFP and performed **aTAG-2** dose titrations. Notably, FKBP^F36V^-mEGFP was not degraded upon **aTAG-2** treatment (**Fig. S3F**). Nevertheless, **aTAG-2** selectively inhibited proliferation of FKBP^F36V^-mEGFP EWS502 cells relative to parental cells, with potency similar to that observed previously (IC_50_ = 1.38 nM). This differential viable decline was again blocked by AP1867 ethyl amide (10 μM) (**Fig. 2H**). Together, these findings support a model in which the differential cytotoxicity of aTAG-2 is not explained solely by target degradation, but instead is consistent with a RIPTAC mechanism contributing substantially to its activity.

We then measured expression of the EWS/FLI target genes *NR0B1* and *NKX2-2* by RT-qPCR in FKBP^F36V^-mEGFP EWS502 cells following **aTAG-2** treatment. **aTAG-2** induced rapid repression of both targets with kinetics nearly identical to those observed in FKBP^F36V^-HA-EWS/FLI cells (**Fig. 2H**), suggesting that degrader activity does not meaningfully contribute to the transcriptional collapse. To test this directly in the original FKBP^F36V^-HA-EWS/FLI model, we used the proteasome inhibitor MG-132 to suppress the degradation component of **aTAG-2** activity (see later discussion in **Fig. 4E**). Under these conditions, **aTAG-2** still repressed target gene expression at 2 hours, as measured by RT-qPCR (**Fig. S3G**).

Overall, these data indicate that **aTAG-2** acts as a ternary-complex dependent degrader and RIPTAC in the Ewing sarcoma FKBP^F36V^ fusion model, with the RIPTAC contribution being primarily responsible for the gene expression/viability changes.

### aTAG-2 behaves similarly in a translocation renal cell carcinoma FKBP^F36V^ fusion model

Given **aTAG-2** induced transactivation in the IRF1 model but transcriptional collapse in the Ewing sarcoma model, we wanted to see its effects across another model where an oncogenic transcription factor has been FKBP^F36V^-tagged. We focused on a model of translocation renal cell carcinoma (tRCC; UOK109) where the oncogenic fusion (*NONO-TFE3*) had monoallelic C-terminal knock-in of FKBP^F36V^-HA.

Because the knock-in is monoallelic (i.e. an untagged copy of the fusion remains), we expected the cell line to respond to the gain-of-function molecule **aTAG-2** (RIPTAC-like) more robustly than dTAG which induces a loss-of-function phenotype (protein degradation). dTAG-V1 was able to selectively inhibit proliferation of NONO-TFE3-FKBP^F36V^-HA UOK109 cells compared to parental with a large therapeutic window (day 12 IC_50_ Parental/FKBP^F36V^ = 11.9 μM/683 nM ∼ 17x); however, **aTAG-2** was both more efficacious and had a larger therapeutic window (day 12 IC_50_ Parental/FKBP^F36V^ = 7.3 μM/26.6 nM = 274x) over a 12-day growth assay (**Fig. S4A**). **aTAG-2** also induced degradation of the HA-tagged fusion protein in NONO-TFE3-FKBP^F36V^-HA UOK109 cells after 24 hours (**Fig. S4B**), an effect that could be competed off by excess of either binder (**Fig. S4C**). The degradation was incomplete compared to dTAG (**Fig. S4B**), providing further evidence that degradation alone is not responsible for the activity of **aTAG-2**.

RT-qPCR analysis of TFE3 fusion targets *TRIM63* and *ANGPTL2* revealed a biphasic response. Initially, **aTAG-2** (50 nM) modestly induced expression of *TRIM63* and *ANGPTL2* (0-30 minutes). Following this, **aTAG-2** robustly repressed both targets in a time-dependent manner, with peak activity around 16-24 hours. Therefore, in the context of a tRCC cell line with a FKBP^F36V^-tagged fusion, **aTAG-2** robustly inhibits cell proliferation, initially transactivating TFE3 fusion targets then repressing them, while also degrading the TFE3 fusion. These data indicate that this paradoxical behavior, degradation and transcriptional repression caused by a putative small molecule transcriptional activator, occurs in multiple fusion oncogene models.

### aTAG-2 erodes chromatin accessibility at p300 and EWS/FLI targets

Having established that the paradoxical transcriptional effects of **aTAG-2** were not restricted to the EWS-FLI context, we sought to gain more insight into the chromatin alterations being induced by drug treatment. To do so, we performed ATAC-seq following 16-hour treatment of FKBP^F36V^-HA-EWS/FLI EWS502 cells with either DMSO, 5 nM **aTAG-2**, or combination of 5 nM GNE-781 + AP1867 ethyl amide (binders). Manual inspection of high confidence EWS/FLI targets (*NR0B1, FCGRT, CCND1, GSTM4*) revealed loss of chromatin accessibility specifically in **aTAG-2** treated cells (**Fig. 3A**). Motif analysis on consensus **aTAG-2** lost ATAC-seq peaks (lost in both **aTAG-2** vs. DMSO and vs. binders comparisons) revealed strong enrichment of the EWS/FLI consensus sequence (rank 1, *P*=1e-568) (**Fig. 3B**). There was also significant enrichment of long GGAA-microsatellite repeats at EWS/FLI-bound sites that lost ATAC-seq signal vs. those with stable ATAC-seq signal (*P*=6.8e-42, OR = 5.42 by chi-squared test, **Fig. S5A**), consistent with preferential collapse of the most EWS/FLI-dependent sites first.

**Fig. 3.**
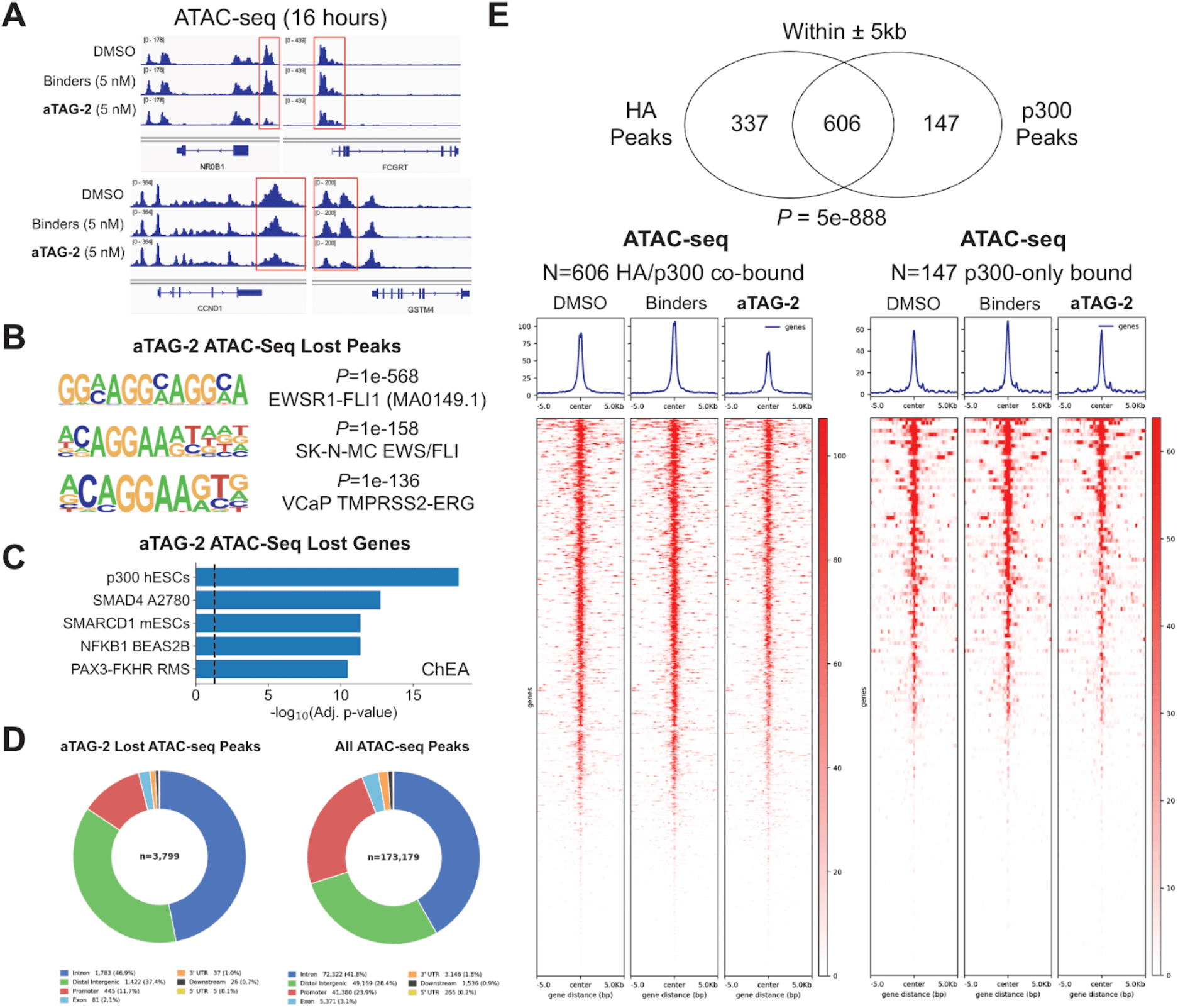
aTAG-2 decreases chromatin accessibility at p300 and EWS/FLI targets. A) FKBP^F36V^-HA-EWS/FLI EWS502 cells treated with DMSO, aTAG-2, or binders (mAP1867-CONHEt + GNE-781) at 5 nM for 16 hours and profiled by ATAC-seq. Chromatin accessibility changes at selected EWS/FLI targets are shown (*NR0B1, CCND1, FCGRT, GSTM4*). B) HOMER motif analysis of consensus ATAC-seq peaks lost by aTAG-2 (lost in aTAG-2 vs. DMSO and aTAG-2 vs. binders comparisons). C) Enrichr gene ontology analysis on unique genes corresponding to consensus aTAG-2 lost ATAC-seq peaks (ChEA gene set database). D) Genomic location of consensus aTAG-2 lost ATAC-seq peaks or all ATAC-seq peaks (see **Methods**) as reference. E) Overlap of FKBP^F36V^-HA-EWS/FLI peaks with p300 peaks (MACS2-based peak calling) in DMSO-treated FKBP^F36V^-HA-EWS/FLI EWS502 cells. *P*-value calculated by hypergeometric test. ATAC-seq signal heatmaps for HA/p300 co-bound peaks and p300-only bound peaks are shown. HA-only bound peaks are shown in Fig. S5D.

**Fig. 4.**
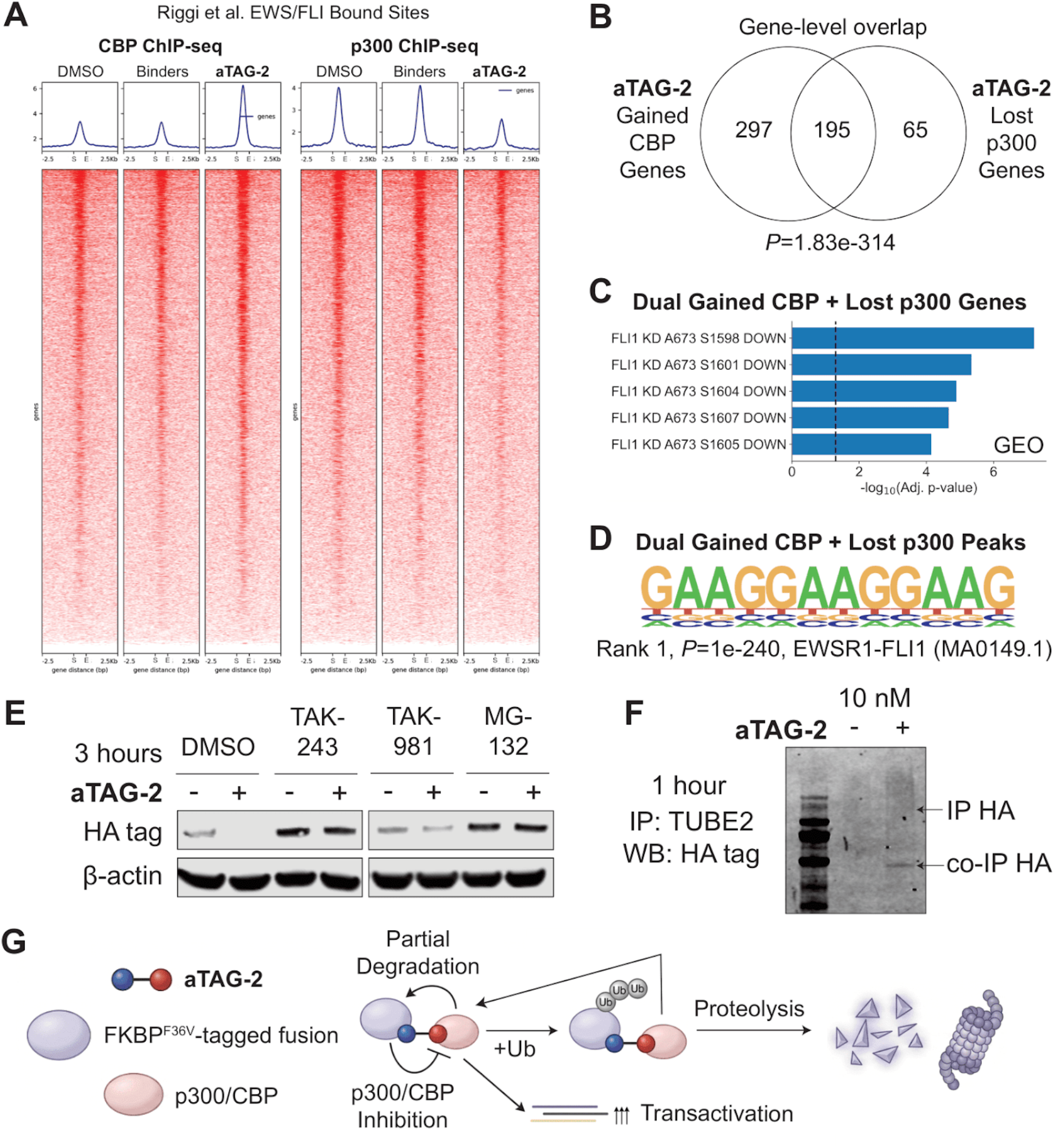
aTAG-2 replaces p300 with CBP at EWS/FLI bound sites and degrades FKBP^F36V^-EWS/FLI in a E1/proteasome-dependent manner. A) FKBP^F36V^-HA-EWS/FLI EWS502 cells treated with DMSO, aTAG-2, or binders (mAP1867-CONHEt + GNE-781) at 10 nM for 16 hours were profiled by CBP, p300, and HA ChIP-seq. P300 and CBP occupancy heatmaps at Riggi et al. EWS/FLI bound sites^85^ are shown. B) Overlap between unique genes with consensus CBP ChIP-seq peaks gained by aTAG-2 (gained both vs. DMSO and vs. binders), and consensus p300 ChIP-seq peaks lost by aTAG-2. *P*-value calculated by hypergeometric test. C-D) Enrichr gene ontology analysis (C) and Homer motif analysis (D) of dual CBP-gained and p300-lost peaks/unique genes. E) Western blot for HA-tagged EWS/FLI in FKBP^F36V^-HA-EWS/FLI EWS502 cells treated with aTAG-2 at 10 nM for 3 hours in the presence of inhibitors (1 μM, except MG-132 at 10 μM; TAK-243: UBA1 inhibitor, TAK-981: SUMOylation inhibitor, MG-132: proteasome inhibitor, lanes are from the same blot & exposure). F) Western blot for HA-tagged EWS/FLI in FKBP^F36V^-HA-EWS/FLI EWS502 cells treated with aTAG-2 at 10 nM for 1 hour following TUBE2 (ubiquitin) pulldown, in the presence of MG-132 (10 μM). Total ubiquitin loading control is shown in Fig. S7C. G) Model for aTAG-2 mechanism of action, where it can induce transactivation, partial target degradation, and inhibit p300/CBP through a RIPTAC-like mechanism depending on context.

Gene ontology analysis of the unique genes corresponding to the consensus **aTAG-2** lost ATAC-seq peaks identified significant enrichment of a p300 hESC target gene set (adjusted *P*=7.24e-19) as well as EWS/FLI target genes (adjusted *P*=5.11e-3) (**Fig. 3C, Fig. S5B**). Consensus **aTAG-2** gained ATAC-seq peaks also displayed enrichment at genes inhibited by EWS/FLI (adjusted *P*=5.77e-5, **Fig. S5B-C**). Lost peaks were heavily biased towards enhancers/intergenic regions (84.3% of peaks) with significant enrichment relative to baseline (i.e. all unique ATAC-seq peaks, OR = 2.3, *P*=2.6e-90 by Fisher exact test). In addition, significantly less promoter-annotated peaks were present among the consensus **aTAG-2** lost ATAC-seq peaks compared to baseline (OR = 0.42, *P*=6.1e-79 by Fisher exact test) (**Fig. 3D**), consistent with enhancer/co-activator collapse.

We next performed p300 and HA ChIP-seq in DMSO-treated FKBP^F36V^-HA-EWS/FLI EWS502 cells to determine the genomic occupancy of p300 and EWS/FLI at baseline. There was very strong overlap between HA and p300 peaks (*P*=5e-888 by hypergeometric test, **Fig. 3E**). **aTAG-2** decreased chromatin accessibility at HA/p300 co-bound peaks, but failed to do so for peaks bound by p300 alone (**Fig. 3E, Fig. S5D**). Therefore, the chromatin effects of **aTAG-2** are concentrated at FKBP^F36V^-HA-EWS/FLI-bound loci, consistent with the RNA-seq and proximity-driven inhibition or sequestration of p300/CBP in *cis* rather than global loss of p300 function.

### aTAG-2 replaces p300 with CBP on chromatin

We next generated ChIP-seq for p300, CBP, and HA following DMSO, **aTAG-2**, or binder treatment (10 nM, 16 hours) in FKBP^F36V^-HA-EWS/FLI EWS502 cells. To assess ternary complex formation on chromatin, we analyzed changes in CBP and p300 occupancy at high confidence sites bound by EWS/FLI.^85^ As expected, CBP occupancy increased significantly at these sites when cells were treated with **aTAG-2**. To our surprise however, p300 occupancy actually decreased at these sites (**Fig. 4A, Fig. S6A**). We envisioned a model where **aTAG-2** was causing p300 to be replaced by CBP on chromatin.

To test this, we analyzed the unique genes corresponding to consensus CBP peaks gained by **aTAG-2** treatment and consensus p300 peaks lost by **aTAG-2** treatment. There was an extremely strong overlap, with 75% of genes losing p300 also gaining CBP (*P*=1.8e-314 by hypergeometric test) (**Fig. 4B**). These dual p300-lost, CBP-gained genes strongly enriched for EWS/FLI targets including *NR0B1* (**Fig. 4C, Fig. S6B**), as well as the EWS/FLI transcription factor binding motif (*P*=1e-240) (**Fig. 4D**). Despite being equipotent for CBP/p300 ternary complex formation (**Fig. 1F-G**), **aTAG-2** exchanged p300 for CBP on chromatin in FKBP^F36V^-HA-EWS/FLI EWS502 cells. This observation may reflect the known high baseline p300 co-occupancy with EWS/FLI, a saturated state in which **aTAG-2** cannot further recruit p300 but can substitute CBP in its place.^82,83^

### aTAG-2 acts as a proteasome and ubiquitination-dependent degrader of FKBP^F36V^-HA-EWS/FLI

We lastly analyzed HA ChIP-seq in **aTAG-2** treated cells. Consistent with the degrader activity we previously observed, **aTAG-2** caused ∼2-fold loss of HA peaks relative to DMSO- or binder-treated cells (**Fig. S7A**). These lost peaks were enriched at EWS/FLI target genes (adjusted *P*=1.59e-11, **Fig. S7B**).

To further characterize the mechanism of targeted protein degradation we profiled **aTAG-2** in the presence of various inhibitors including TAK-243 (E1 inhibitor), TAK-981 (SUMOylation inhibitor), and MG-132 (proteasome inhibitor) after 3 hours. The degradation of FKBP^F36V^-HA-EWS/FLI was dependent on E1 activity (TAK-243) and the proteasome (MG-132) (**Fig. 4E**). To confirm ubiquitination, we performed tandem ubiquitin binding entity 2 (TUBE2) pulldown of **aTAG-2** treated cells and observed the appearance of a HA-ubiquitin smear. A substantial amount of non-ubiquitinated HA was also pulled down, likely due to induced ubiquitination of EWS/FLI interactors resulting in co-IP (**Fig. 4F, Fig. S7C**). The degradation of FKBP^F36V^-HA-EWS/FLI was independent of p300/CBP acetyltransferase activity (using inhibitor A-485) and TRIM8 activity (the primary E3 ligase for EWS/FLI, tested using dox-inducible sgRNAs) (**Fig. S7D-E**).

Overall, these results support a model where **aTAG-2** is a proximity-inducing molecule with activity as a transactivator, RIPTAC, degrader, and relocalizer on chromatin. It transactivates tagged IRF1 in a U2OS reporter line and has biphasic activity in tagged NONO-TFE3 tRCC, initially transactivating the fusion before inhibiting target gene expression. In tagged EWS/FLI Ewing sarcoma cells, its dominant effect is rapid transcriptional collapse at fusion-bound loci, consistent with co-activator redistribution/inhibition in *cis*, while degradation proceeds via an E1- and proteasome-dependent pathway. The RIPTAC-like effect appears responsible for the gene expression/viability phenotypes (**Fig. 4G**).

## Discussion

Here we developed a modular platform to test whether bifunctional small molecules can activate transcription by recruiting endogenous activating machinery to a FKBP^F36V^-tagged transcription factor. Across a panel of candidate aTAG compounds, only a subset produced robust activation of the IRF1 reporter, consistent with the idea that simply recruiting proteins with activating functions is not always sufficient to generate productive transcription in a given configuration. This outcome likely reflects geometric and contextual constraints on induced proximity, including the orientation of the recruited effector relative to the transcription factor and the surrounding transcriptional machinery, which may depend on the position of the FKBP^F36V^ tag and the ability of recruited factors to engage multi-protein complexes. In contrast, **aTAG-2** (mAP1867-C8-GNE781) was a clear exception in this system, producing potent reporter activation and displaying classic bifunctional behavior (hook effect, inactivity of monovalent ligands, and competition by excess binders), supported by NanoBiT evidence for p300/CBP-FKBP^F36V^ ternary complex formation.

Unexpectedly, the functional output of **aTAG-2** was strongly context dependent. In a Ewing sarcoma model expressing FKBP^F36V^-tagged EWS/FLI, **aTAG-2** caused rapid repression of EWS/FLI target genes and selective loss of viability, with repression kinetics (≤ 1-2 h) faster than would be expected from indirect stress responses. This divergence suggests that whether proximity produces activation vs. collapse likely depends on the baseline epigenetic state of the targeted program. IRF1 loci may have headroom for added coactivator recruitment, whereas EWS/FLI-driven enhancers, already heavily coactivator-occupied, may be vulnerable to forced rewiring of coactivator dynamics. Concretely, **aTAG-2** seems to be replacing active p300 recruited by the native EWS TAD to EWS/FLI motifs with a mixture of CBP and p300 that are simultaneously inhibited by the recruiting compound. The net result of this action is repression in *cis*. Other compounds including **aTAG-1** and **aTAG-6** displayed similar repressive behavior in the tagged Ewing sarcoma cell line, suggesting that TOVER in oncogenic fusion contexts may be challenging and require novel non-inhibitory ligands of activating epigenetic machinery.

Chromatin profiling supports a *cis*-localized coactivator collapse model: **aTAG-2** preferentially eroded accessibility at EWS/FLI-bound sites, enriched for EWS/FLI motifs, while sparing p300-only sites, consistent with local disruption rather than global p300 shutdown. Strikingly, ChIP-seq revealed increased CBP occupancy but decreased p300 occupancy at fusion-bound loci, indicating that **aTAG-2** can drive paralog exchange on chromatin despite similar p300/CBP ternary complex formation potency. This provides a concrete mechanism by which an “activator recruitment” strategy can become repressive in a saturated enhancer environment.

Mechanistically, our data indicate that **aTAG-2** is pleiotropic, exhibiting degrader- and RIPTAC-like behaviors. **aTAG-2** partially degraded FKBP^F36V^-HA-EWS/FLI in a ternary-complex dependent manner and via an E1- and proteasome-dependent pathway. However, comparable viability decline and rapid repression in cells expressing an unrelated FKBP^F36V^-mEGFP fusion, without detectable degradation of the tagged protein, argues that RIPTAC-driven functional disruption is a major contributor to the transcriptional phenotype, with degradation acting as an additional output rather than the sole driver.

Overall, these results demonstrate that bifunctional compounds may often behave in more complex ways than we currently appreciate. In this case, a ligand previously shown to be a transcriptional activator can produce multiple activities that vary by time and cellular context, including activation, relocalization on chromatin, RIPTAC-like inhibition, and degradation. For coactivators such as p300/CBP, large scaffolding proteins with many interaction surfaces, forced proximity may inhibit more functions than monovalent bromodomain or catalytic inhibitors due to steric blockade, helping explain the efficacy of **aTAG-2** in fusion-driven settings.

More broadly, these results emphasize that induced proximity with a given ligand does not encode a fixed functional outcome. Instead, whether recruitment produces activation or collapse is dictated by the pre-existing regulatory state (e.g. coactivator occupancy) so binary binding and even ternary complex formation are insufficient to predict directionality. As more complex proximity inducing perturbagens are discovered, we anticipate that these compounds will frequently employ multiple context-dependent mechanisms of action.

## Acknowledgements

This work was supported by the Hertz Foundation Fellowship (A.Sa.), Herchel Smith Graduate Fellowship (A.Sa.), F32 CA284750 (M.J.B.), T32 HL007574 (C.N.W.), AACR-Exelixis Renal Cell Carcinoma Research Fellowship (C.N.W.), NIH R01CA262188 (E.S.F.), NCI R35CA283977 (K.S.), NCI R01CA286652 and R01CA279044 (S.R.V.), NIH R01HL082945 and P01CA066996 (B.L.E.), the Howard Hughes Medical Institute (B.L.E.), the Edward P. Evans Foundation (B.L.E.), the Adelson Medical Research Foundation (B.L.E.), the Break Through Cancer foundation (B.L.E.), the Svenson Fellowship (W.J.G.), the Lubin Scholar Award (W.J.G.), the Chleck Family Foundation (W.J.G), and the Briger Foundation for Oncology Research Award (W.J.G).

## Author Contributions

A.Sa. conceptualized this study and did the majority of experiments including chemical synthesis of aTAGs, establishing the reporter, profiling in cellular models, RT-qPCR, Western blots, RNA/ChIP/ATAC-seq, analysis of the RNA/ChIP/ATAC data, and wrote the manuscript. M.C. assisted with Western blots and cloning. E.J.Z. assisted with chemical synthesis. S.S. performed Western blots and some viability experiments in the tRCC model. C.N.W. established the tagged translocation renal cell carcinoma model. M.J.B. assisted with study design and established the dox-inducible TRIM8 KO line. A.So. assisted with cloning plasmids. K.A.D. and J.K.R. performed global proteomics experiments and analysis. E.F., K.S., S.R.V., B.L.E., and W.J.G. provided funding and supervised study design. W.J.G. conceptualized this study, provided guidance on experimental design and edited the manuscript.

## Methods

### Cell Culture

293T and U2OS cells were obtained from ATCC. UOK109 NONO-TFE3-FKBP^F36V^ cells were generated in Dr. Srini Viswanathan’s laboratory (DFCI), and EWS502 FKBP^F36V^-EWSR1-FLI1 cells were generated in Dr. Kim Stegmaier’s laboratory.^34^ 293T, U2OS, and UOK109 cells were cultured in DMEM supplemented with 10% FBS, whereas EWS502 cells were cultured in RPMI supplemented with 15% FBS. All cell lines were maintained in media supplemented with 100 IU/mL penicillin and 100 μg/mL streptomycin and grown at 37°C in 5% CO_2_.

### Plasmids

The U2OS IRF1 reporter was generated from the pLminP Luc2P backbone (see Addgene #90363),^86^ where destabilized firefly luciferase-hPEST is under control of minP with upstream consensus transcription factor binding sites in a lentiviral vector that expresses EmGFP from a different promoter. We ordered an insert from Twist Bioscience containing 4 copies of the IRF1 consensus sequence (GAAAGTGAAAGTGAAAGT) that we Gibson cloned (NEB HiFi DNA assembly) into this backbone in place of the existing TF binding sites. IRF1-FKBP^F36V^-mCherry and FKBP^F36V^-mEGFP were ordered from Twist as a codon optimized entry vectors (pTwist-ENTR) that were Gateway cloned (Invitrogen LR clonase II) into either lentiviral dox-inducible expression vector pLIX403 (Addgene #41395) or lentiviral constitutive expression vector pLX311 (Broad Institute). NanoBiT plasmids (FKBP^F36V^-LgBiT, CBP(BD)-SmBiT, p300(BD)-SmBiT) were ordered from Twist as codon optimized pCMV expression vectors (pTwist-CMV). Sequences were verified through PlasmidSaurus whole-plasmid sequencing.

### Transient Transfection and Stable Cell Line Creation

Plasmids were transfected into 293T cells using TransIT-LT1 transfection reagent (1:3 ratio) following the manufacturer’s protocol. Lentivirus was generated transfecting psPAX2 (Addgene: 12260), pMD2.G (Addgene: 12259), and the cloned lentiviral plasmid (1:1:1 ratio) into 293T cells. Lentivirus was collected 2 days after transfection and stable cell lines were established infecting with filtered (0.45 μm filter) lentivirus and polybrene (10 μg/mL). Cells were switched to selection media (U2OS: 5 μg/mL puromycin, EWS502: 10 μg/mL blasticidin) 2 days after infection.

### U2OS Reporter Assays

U2OS reporter cells were resuspended in complete growth media (2,000 cells/50 μL) and doxycycline was added (500 ng/mL). The resulting mixture was plated in white 384-well plates with varying concentrations of compounds (plated with compound printer) using a total volume of 50 μL/well. Firefly luciferase activity (IRF1 reporter activity) was assessed with reconstituted OneGlo reagent (10 μL/well), which was mixed on a Multidrop Combi for 1 minute prior to the reading on a EnVision 2105 multimode plate reader. Luciferase activity was readout 48 hours after compound addition unless otherwise stated.

### NanoBiT Assay

293T cells were seeded in 6-well plates and co-transfected with FKBP^F36V^-LgBiT and SmBiT constructs 24 hours after plating (1 μg plasmid each, 6 μL TransIT-LT1). After one day, the cells were passaged into white 384-well plates (10,000 cells/well) containing varying concentrations of compound (plated with a compound printer). One day after treatment, fluorofurimazine (Ambeed, diluted to final concentration of 40 μM) was added to the cells, mixed, and luminescence was monitored using an EnVision 2105 multimode plate reader.

### Cell Viability Assays

For 4-day growth assays, cells were resuspended in complete growth media (EWS502: 1,500 cells/50 μL, UOK109: 1,000 cells/50 μL) and plated in white 384-well plates with varying concentrations of compounds (plated with compound printer) using a total volume of 50 μL/well. Viability was assessed using reconstituted cell titer glo (CTG) reagent (20 μL/well), which was mixed, left to stand for 10 minutes in the dark, and readout using an EnVision 2105 multimode plate reader. Viability was readout 96 hours after compound addition unless otherwise stated.

### Confluence Assays

For 12-day growth curves, NONO-TFE3-FKBPF36V-HA and parental UOK109 cells were plated in black, clear-bottom 96-well plates (Corning, Ref. 3603) at 500 cells per well in a total volume of 100 μL. Compounds were dispensed using a compound printer (Tecan D300e) at the indicated concentrations at the time of plating. Media and compounds were refreshed every 4 days. Plates were imaged daily in a Celigo Image Cytometer after establishing appropriate focus levels for each plate for each day (well mask 80%, confluence 1).

### Immunoblotting and Degradation Assays

Cells were split into 6-well plates (300k cells/well) and grown overnight. Cells were treated with compounds at the indicated concentration for the indicated amount of time then trypsinized, pelleted, flash frozen on liquid nitrogen, and stored at -80C. The pellets were thawed on ice, and lysed in RIPA buffer (typically 30 µL) with protease inhibitors (cOmplete, Mini Protease Inhibitor Cocktail or Halt Protease Inhibitor Cocktail, EDTA-Free) for 30 min. Lysates were clarified (21,000g, 10 min, 4C) and protein concentration was determined by BCA assay. Samples were normalized to 15 µg protein in 20 µL of loading buffer (5 µL of 4X LDS buffer + DTT [50 mM]), and denatured on a heat block at 95C for 5 mins. Samples were resolved on 4-12% NuPAGE Bis-Tris gels (120V) and transferred to PVDF membranes using the iBlot system (7 min, P0). Membranes were blocked in Intercept blocking buffer (1 h, RT), incubated overnight at 4 °C with primary antibodies (for UOK109 blots: β-actin 1:3000 [Mouse mAb 8H10D10 3700S]; HA 1:1000 [Mouse mAb 6E2 2367S]; TFE3 1:1000 [ZoomAb Rabbit mAb Clone 1G5 2RB1272-25ul]; for EWS502 blots: β-actin 1:5000 [Mouse mAb 8H10D10 3700S], HA 1:1000 [Rabbit mAb C29F4 3724S], GFP 1:1000 [Mouse mAb 4B10 2955S], TRIM8 1:1000 [Rabbit mAb F3C3T 85434S], Ubiquitin 1:1000 [Mouse mAb FK2 39269S]), washed in TBS-T (x3), then incubated with IRDye 680LT anti-mouse and IRDye 800CW anti-rabbit secondaries (1:10000, 1 h, RT). Membranes were washed with TBS-T (x3) and imaged on a near-infrared imager.

### TUBE2 Pulldown Assay

EWS502 FKBP^F36V^-EWSR1-FLI1 were split into 10 cm dishes (5M cells/plate) and grown overnight. All plates were treated with 10 μM MG-132, and then treated with either DMSO or aTAG-2 (10 nM). After 1 hour, cells were trypsinized and pelleted. Cells were resuspended in lysis buffer containing 1% NP-40, 150 mM NaCl, 20 mM Tris pH 7.5, 100 μM PR-619, 5 mM N-ethylmaleimide (NEM), 5 mM o-phenanthroline, 1X Halt protease inhibitor cocktail, and 20 μM MG-132. Cells were lysed for 30 minutes at 4C and lysates were clarified (21,000g, 10 min, 4C). TUBE2 high capacity magnetic beads (UM502M, 100 μL/pulldown) were washed twice with TBS-T and allowed to bind clarified lysate for 3 hours at 4C with rotation. The beads were washed three times with TBS-T containing 100 μM PR-619, 5 mM NEM, 5 mM o-phenanthroline, 1X Halt protease inhibitor cocktail, and 20 μM MG-132. Protein was eluted off beads using 1.5X LDS buffer containing 50 mM DTT, heating at 95C for 10 minutes. The beads were centrifuged for 1 minute at 21,000g, placed on a magnetic rack, and the lysate was loaded onto a 4-12% NuPAGE Bis-Tris gel for a HA Western blot.

#### RT-qPCR

Cells were split into 12-well plates (150k cells/well) and grown overnight. Cells were treated with compound at the indicated concentration for the indicated time, and then trypsinized and pelleted. Cells were lysed in RLT buffer (350 μL) by hard vortexing for 30 seconds and RNA was purified using the RNAeasy Mini Kit. 1 μg of total RNA was reverse transcribed using SuperScript IV VILO Master Mix (10 μL total volume) at 50C for 20 mins, followed by heat denaturation at 85C for 5 mins. The resulting cDNA was diluted 1:12 with ddH_2_O. All qPCR reactions were performed using SYBR Green Master Mix (10 μL total volume, 2 μL diluted cDNA, 5 μL Master Mix, 300 nM primer concentration). The amplification was performed in hard-shell 384-well PCR plates, thin wall, skirted, clear/white (BioRad), sealed with an adhesive cover. The QuantStudio 6 Flex Real-Time PCR machine and the accompanying QuantStudio Real-Time PCR software v.1.7 (Thermo Fisher Scientific) was used to produce and analyze data. The delta-threshold cycle number (ΔCt) was calculated as the difference in threshold cycle number (Ct) between the gene of interest and *GAPDH*. The ΔΔCt was calculated as the difference between the ΔCt of a particular sample and the average ΔCt of the DMSO-treated samples. The fold change in gene expression was calculated by using the following exponential equation: fold change in expression of treatment vs DMSO = 2^−ΔΔCt^. qPCR primers used in this study: NKX2-2_F: GTCAGGGACGGCAAACCAT, NKX2-2_R: GCGCTGTAGGCAGAAAAGG, NR0B1_F: AGCACAAATCAAGCGCAG, NR0B1_R: GAAGCGCAGCGTCTTCAA, TRIM63_F: ATGAATGAGAGGCCCCCAGAT, TRIM63_R: TACCCTAGTCCCTGCTCTCTG, ANGPTL2_F: GAACCGAGTGCATAAGCAGGA, ANGPTL2_R: GTGACCCGCGAGTTCATGTT, GAPDH_F: GTCTCCTCTGACTTCAACAGCG, GAPDH_R: ACCACCCTGTTGCTGTAGCCAA.

### RNA-seq

EWS502 cells were split into 12-well plates (150k cells/well) and treated with compound (batch 1: DMSO, 5 nM aTAG-2, 5 nM GNE781 + mAP1867-CONHEt; batch 2: DMSO, 500 nM aTAG-6; batch 3: DMSO, 250 nM aTAG-1 in duplicate) for 16 hours. Cells were trypsinized and pelleted. For batch 1, TRIzol (Invitrogen) was added to cells, and following the manufacturer’s protocol, RNA was extracted. RNA concentration was monitored using a Qubit Fluorometer (ThermoFisher), and RNA integrity was analyzed using an Agilent Bioanalyzer. The NEBNext Ultra II RNA Library Prep Kit (Illumina) was used to prepare a RNA-seq library. A NovaSeq 6000 machine (Illumina) was used for paired-end 150 bp RNA-sequencing. STAR/RSEM^87,88^ was used to align RNA-seq reads to the GENCODE v38 transcript reference^89^ and generate a count matrix. DESeq2^90^ was used to calculate log_2_(fold-changes) and adjusted *P*-values. For batches 2-3, 100 μL of DNA/RNA Shield (Zymo) was added to cells, and the cells were lysed by hard vortexing for 30 seconds. The solution was sent to PlasmidSaurus for 3’ Tag-seq and default analysis to generate log_2_(fold-changes)/P-values. A pre-ranked GSEA^91^ was performed using either the DESeq2 *t*-statistic (batch 1) or the PlasmidSaurus signed log_10_(FDR) (batch 2-3) as a rank metric. Kinsey et al. Ewing sarcoma target gene sets were used for this analysis.^84^

### ATAC-seq

EWS502 FKBP^F36V^-EWSR1-FLI1 cells were passaged into 6-well plates (500k cells/well) and treated with DMSO, 5 nM aTAG-2, or 5 nM GNE781 + mAP1867-CONHEt for 16 hours. Cells were washed once with cold PBS and cryopreserved in 10% DMSO, 40% complete growth media, and 50% FBS. Cryopreserved cells were shipped on dry ice to Novogene for ATAC-seq library preparation and sequencing. In brief, nuclei were extracted from samples, undergo quality control, resuspended in Tn5 transposase reaction mix (including adapters) and incubated for 30 mins at 37°C. PCR was performed to amplify the library which was then purified using AMPure beads. Library quality was assessed on a Tapestation 4150 and it is diluted to 2-4 ng/μL. The insertion size of the library was detected by NGS3K and the concentration was quantified and normalized using qPCR. Vazyme Hyperactive ATAC-Seq library prep kit (Illumina) was used for library construction. Sequencing was performed on a NovaSeq X Plus (Illumina). During analysis, adaptors were trimmed with fastp^92^ and reads were aligned using bwa mem^93^ to hg38. Mitochondrial reads were discarded with samtools^94^ and duplicates were removed with Picard. Reads were restricted to those that were properly paired and mapped (both itself and its mate), passing QC, and having mapping quality of at least 30. Bam files were converted to RPGC normalized (effective genome size: 2913022398) bigwig files using bamCoverage (Deeptools^95^) for visualization in IGV.^96^ Differential binding analysis was performed using csaw.^97^ Genome-wide accessibility was quantified by counting ATAC-seq reads in sliding genomic windows (150 bp width, 50 bp spacing) using csaw. To estimate local background for each window, counts were also obtained for a 2 kb neighborhood centered on the window, and windows were retained if they showed >3-fold enrichment over this local background and had at least 10 total counts across samples. Filtered window counts were converted to an edgeR DGEList^98^, normalized by TMM^99^, and modeled with a negative binomial GLM using a design matrix with DMSO/Binders as the reference. Differential accessibility for aTAG-2 vs. DMSO/Binders was assessed using a likelihood-ratio test with a fixed dispersion of 0.05. Adjacent windows were then merged into broader regions (≤100 bp gap, maximum width 5 kb), and window-level statistics were aggregated to region-level significance using combineTests, with the strongest window logFC reported per region. Consensus lost peaks were defined based on regions with lost ATAC-seq signal (*P*<0.05) in both comparisons: aTAG-2 vs. DMSO and aTAG-2 vs. Binders. To analyze the genomic annotation of all ATAC-seq peaks we generated a bed file of all unique (non-overlapping) csaw windows (i.e. those with >3-fold enrichment over local background and at least 10 counts) across any of the treatment conditions from the differential binding analysis. ChIPseeker^100^ was used to annotate windows to genes/peak classes (e.g. intergenic, intron, promoter, etc.). Homer motif analysis^101^ (findMotifsGenome.pl with -size 200) was performed on consensus lost peaks. Enrichr gene ontology analysis^102^ was performed using the unique genes annotated for lost peaks. The overlap of these unique genes with Kinsey et al. Ewing sarcoma target gene sets were also assessed; *P*-values were calculated by hypergeometric test. Deeptools computeMatrix (center referencePoint) on reference bed files and plotHeatmap were used to generate heatmaps.

### ChIP-seq

EWS502 FKBP^F36V^-EWSR1-FLI1 cells were passaged into 10 cm dishes (5M cells/plate/IP) and treated with DMSO, 10 nM aTAG-2, or 10 nM GNE781 + mAP1867-CONHEt for 16 hours. Cells were trypsinized and pelleted. The cells were resuspended in 10 mL of PBS containing 2 mM DSG (disuccinimidyl glutarate) and incubated with agitation for 30 mins. Methanol-free formaldehyde was added to the solution (to final concentration of 1%) and the suspension was incubated for another 10 mins with agitation. Glycine (10X solution, CST) was added to a final concentration of 1X and the mixture was incubated for an additional 5 mins with agitation. The cells were pelleted and washed twice with PBS (10 mL) containing Pierce protease inhibitor tablets. The cells were lysed in ChIP lysis buffer (20 mM Tris pH 7.5, 300 mM NaCl, 2 mM EDTA, 0.5% NP-40, 1% Triton X-100) on ice for 30 min at pelleted at 4C (5000 rpm, 10 min). The cell pellet was resuspended in 1 mL of sonication buffer (0.1% SDS, 0.5% N-lauroylsarcosine, 1% Triton X-100, 10 mM Tris-HCl pH 8, 100 mM NaCl, 1 mM EDTA) and sonicated using the Covaris E220 (5% duty cycle, 140W peak power, 200 cycles/burst, 4C, 25 mins, milliTUBE). The sonication mixture was pelleted (13000 rpm, 10 min, 4C) and the supernatant was quantified, then normalized to the lowest concentration. Dynabeads Protein A (50 μL) were washed 3 times with TBS-T then pre-incubated with 10 μg of antibody (anti-HA tag antibody - ChIP Grade (ab9110), P300 (Bethyl, A300-358A), CBP (Bethyl, A300-362A)) at 4C for 3 hours in TBS-T (500 μL). The bead-antibody complexes were mixed with supernatants at 4C overnight. The beads were washed using the SimpleChIP Plus Chromatin Immunoprecipitation Protocol recommendations; specifically this involves 3 low salt washes, 1 high salt wash, and elution in 1X ChIP elution buffer (150 μL) at 65C for 30 mins. Cross-links were reversed by adding 6 µL 5M NaCl and 2 µL Proteinase K to the eluate and mixing at 65C for 16 hours. The DNA was purified by CST spin column (14209S). Library construction and sequencing was performed at Novogene; Ultra II DNA Library Prep Kit (NEB) was used for library construction. Sequencing was performed on a NovaSeq X Plus (Illumina). During analysis, reads were preprocessed, aligned, and processed into bam/bigwig/bed/narrowPeak files using the default ChiLin pipeline (hg38, MACS2-based peak calling).^103,104^ Bedtools^105^ intersect/window were used to determine peak overlap. Consensus gained/lost peaks were determined using csaw-based analysis (|log_2_(FC)|>0.5), in an identical manner to the ATAC-seq data. Motif analysis and gene ontology were also performed following the ATAC-seq methods. Riggi et al. EWS/FLI bound enhancers^85^ were used as a reference for EWS/FLI binding in this analysis.

### Global Proteomics

EWS502 FKBP^F36V^-EWSR1-FLI1 cells were passaged into 10 cm dishes (5M cells/plate) and grown overnight. Cells were treated with DMSO or aTAG-2 (10 nM) for 3 hours in triplicate. Cells were trypsinized, pelleted, and flash frozen in liquid nitrogen. Global proteomics was performed as previously described.^106^

### Statistical Analysis

Statistical tests (two-sided) were performed with Python. Dose-response data were fit by nonlinear regression (scipy), using a four-parameter sigmoidal model for monotonic dose responses and, for non-monotonic “hooked” curves, a log-concentration-based peaked (log-normal) model with bounded (or variance-weighted) least-squares fitting. For sequencing analyses, multiple-hypothesis testing was controlled using the Benjamini-Hochberg procedure. Hypergeometric and Fisher exact tests were used for peak overlaps and annotation enrichments as described in the Results.

### Chemical synthesis

The synthesis and characterization of the small molecules reported in this paper are described in the **Supplementary Information**.

## Supplementary Figures

**Fig. S1.**
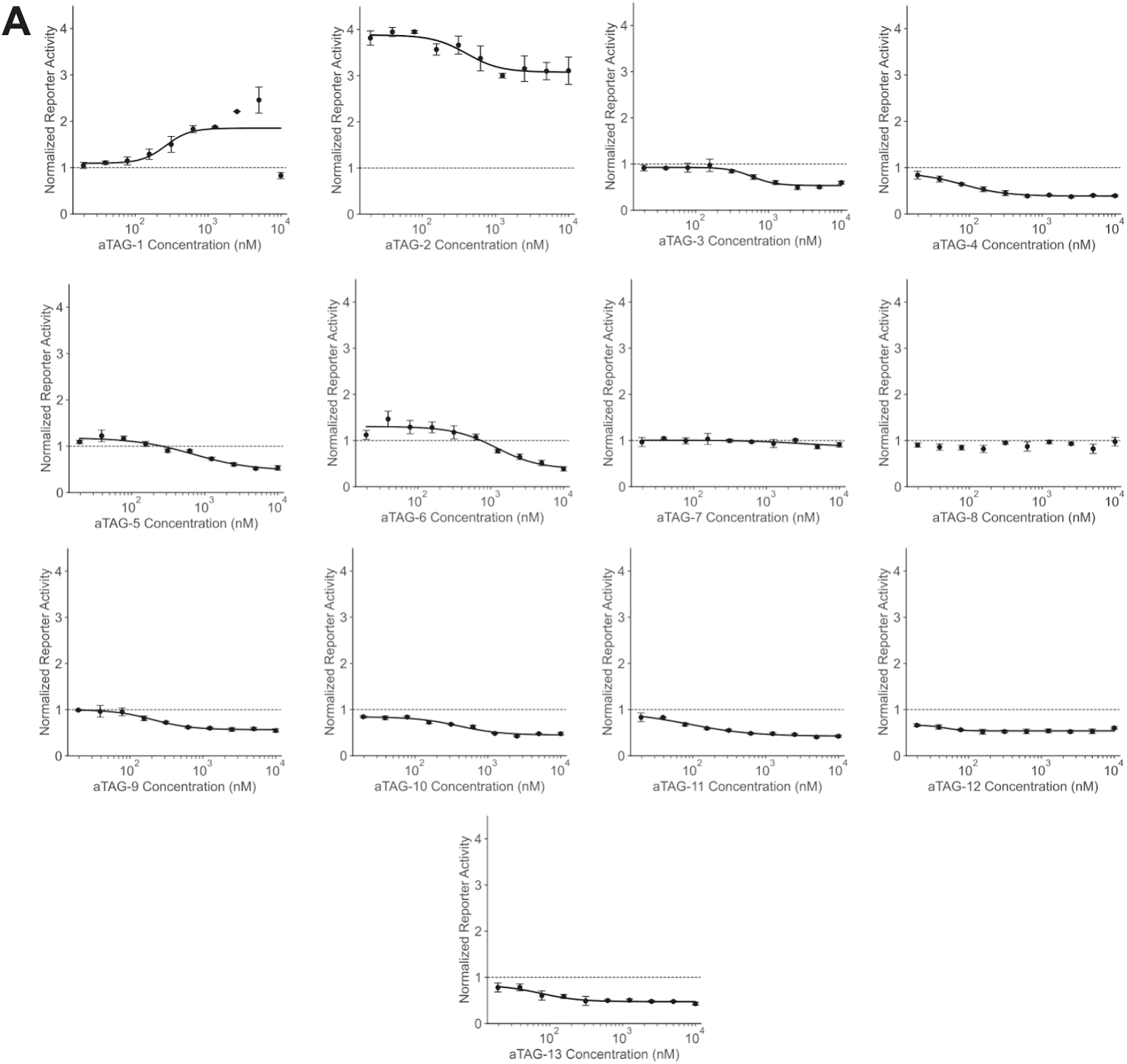
Dose curves of individual aTAG compounds in the IRF1 reporter system. 10-point dose curves of each aTAG compound in the U2OS IRF1 reporter system (normalized FLuc luminescence at 48 hours). Summarized data are shown in Fig. 1A-B.

**Fig. S2.**
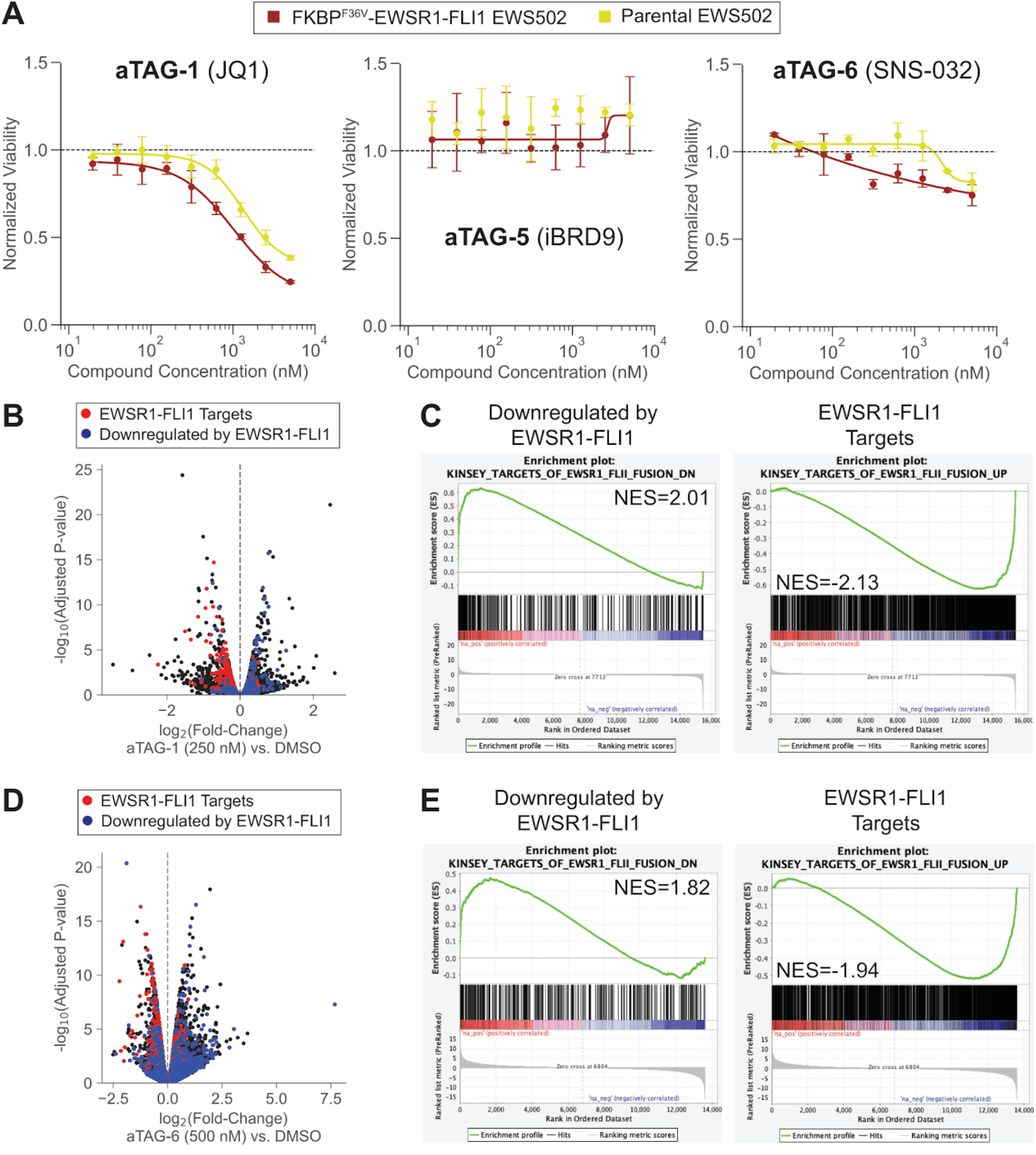
Differential inhibition of FKBP^F36V^-HA-EWS/FLI EWS502 cell growth with other aTAG molecules is accompanied by EWS/FLI target gene repression. A) The impact of other aTAG molecules with transactivating activity in the IRF1 reporter on the differential viability of tagged EWS cells was assessed. Compounds were dose titrated in FKBP^F36V^-HA-EWS/FLI or parental EWS502 cells. Cell viability was read out at 96 hours by CTG. aTAG-1 and aTAG-6 demonstrated differential activity. B,D) RNA-seq volcanoes at 16 hours comparing aTAG-1 (250 nM, B) or aTAG-6 (500 nM, D) to DMSO-treated FKBP^F36V^-HA-EWS/FLI EWS502 cells. Kinsey et al. targets of EWSR1/FLI1 fusion UP are labelled in red (genes transactivated by EWS/FLI), DOWN labelled in blue (genes downregulated by EWS/FLI). C,E) Individual pre-ranked gene set enrichment plots across Kinsey et al. EWS/FLI gene sets from RNA-seq experiment in panel B/D (aTAG-1: panel C, aTAG-6: panel E).

**Fig. S3.**
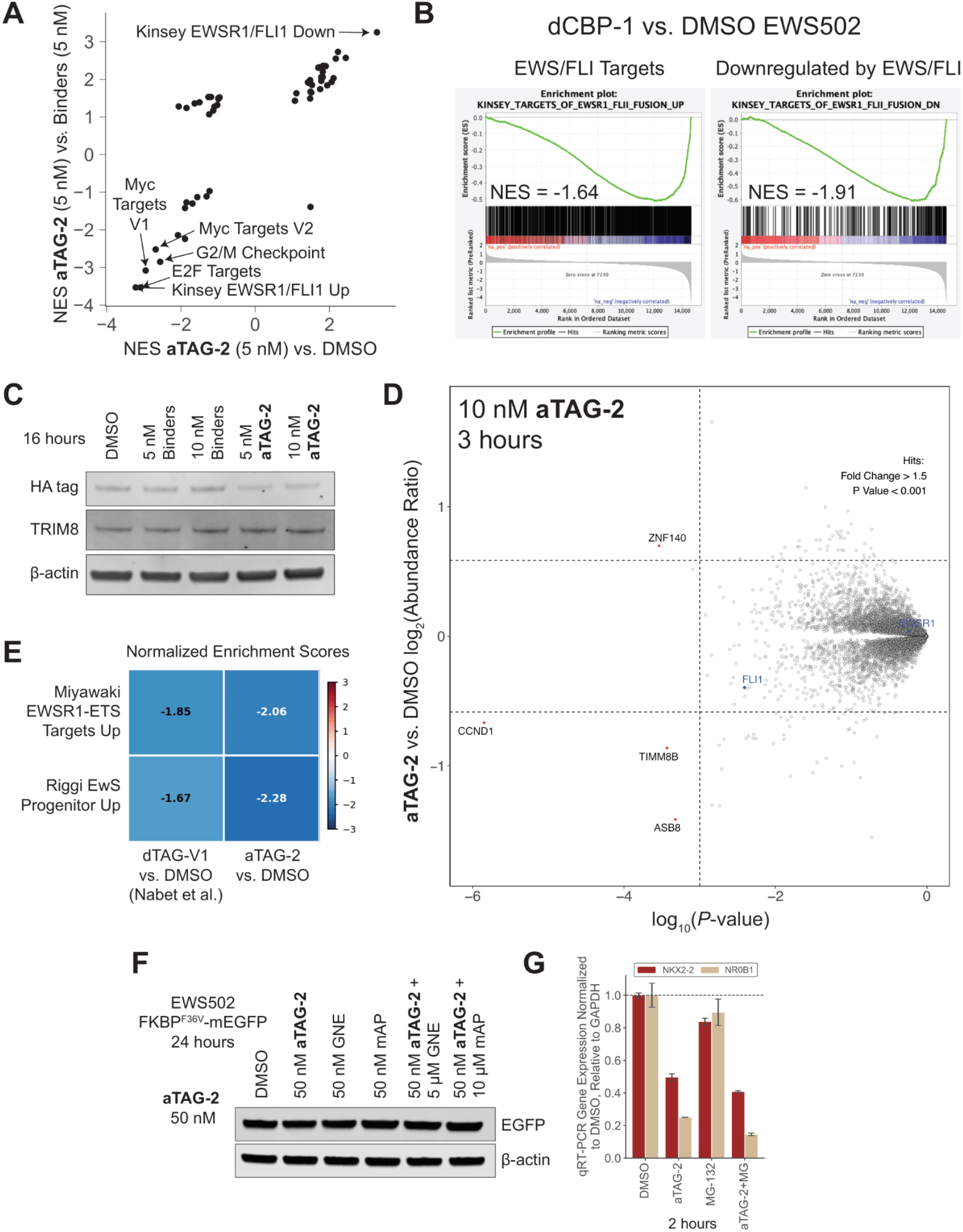
aTAG-2 demonstrates on-target activity. A) Normalized enrichment score (NES) comparison from pre-ranked GSEA across hallmark gene sets (Kinsey et al. EWS/FLI gene sets added) comparing aTAG-2 to DMSO (x-axis) or to binders (y-axis) from RNA-seq experiment in Fig. 2B. B) Pre-ranked GSEA plots across Kinsey et al. EWS/FLI gene sets comparing previously published RNA-seq in dCBP-1 vs. DMSO treated EWS502 cells.^82^ Rank metric was log_2_(fold-change) between the two groups. C) Western blot for HA-tagged EWS/FLI, TRIM8, and actin in FKBP^F36V^-HA-EWS/FLI EWS502 cells treated with aTAG-2 or binders (mAP1867-CONHEt + GNE-781) at the indicated concentrations for 16 hours. D) Global proteomics following aTAG-2 (10 nM) or DMSO treatment of FKBP^F36V^-HA-EWS/FLI EWS502 cells for 3 hours. FLI1 and EWSR1 are labelled. Note: wild-type, untagged EWSR1 is expressed in this cell line confounding readouts of EWSR1 degradation. E) Normalized enrichment score heatmap from pre-ranked GSEA analysis of dTAG-V1 vs. DMSO treated FKBP^F36V^-HA-EWS/FLI EWS502 cells from Nabet at al.^34^ compared to aTAG-2 vs. DMSO treated cells. F) Western blot for EGFP in FKBP^F36V^-mEGFP EWS502 cells treated with aTAG-2 ± competitors or binders. G) RT-qPCR for EWS/FLI targets (NKX2-2, NR0B1; normalized to GAPDH) in the presence of absence of proteasome inhibitor MG-132 (10 μM) and aTAG-2 (10 nM) in FKBP^F36V^-HA-EWS/FLI EWS502 cells at 2 hours.

**Fig. S4.**
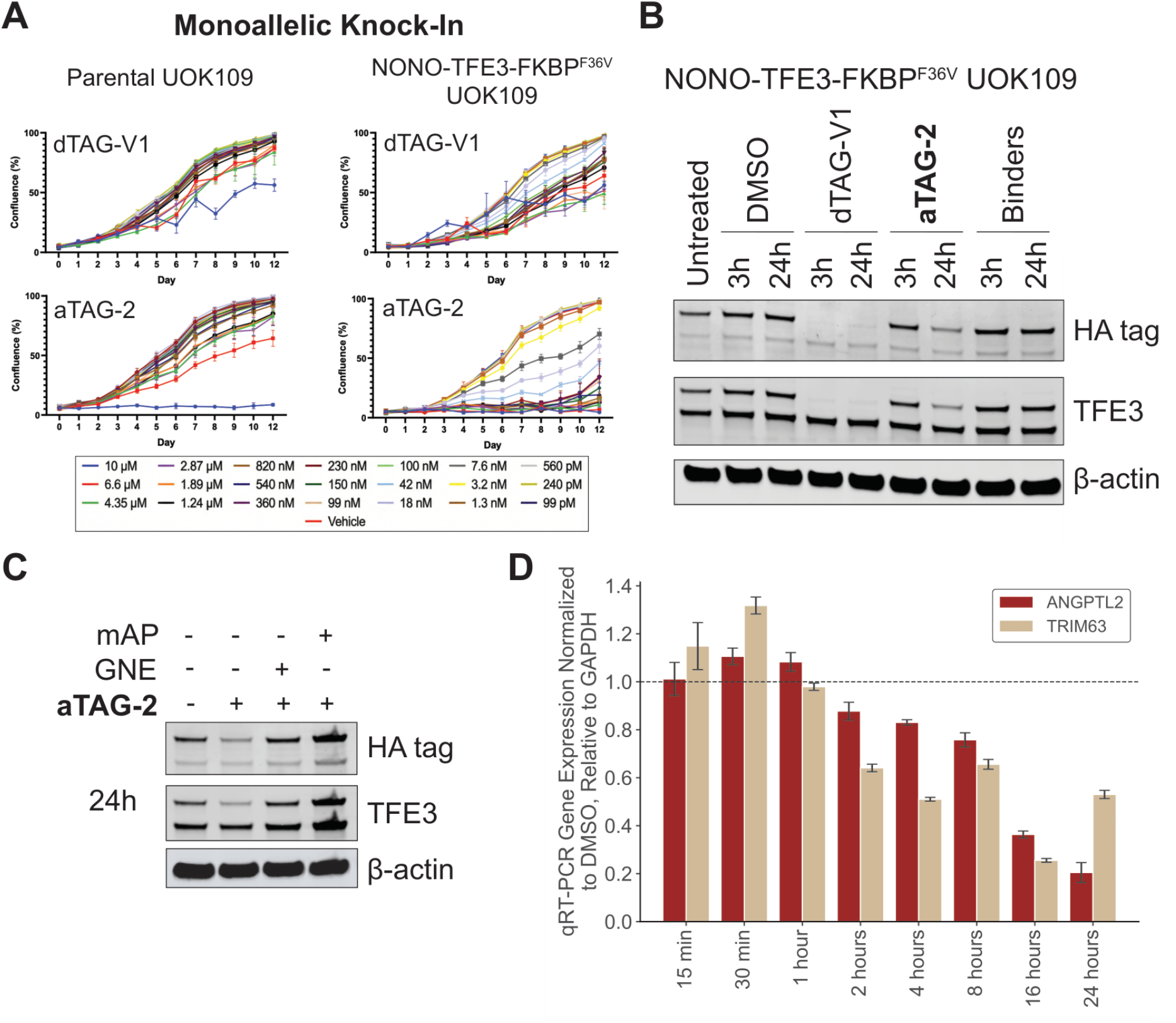
aTAG-2 demonstrates activity in translocation renal cell carcinoma NONO-TFE3-FKBP^F36V^-HA monoallelic knock-in model. A) Dose curves for dTAG-V1 and aTAG-2 in NONO-TFE3-FKBP^F36V^-HA and parental UOK109 tRCC cells. Cell confluence was read out on Celigo daily for 12 days. B) Western blot for HA and TFE3 in NONO-TFE3-FKBP^F36V^-HA UOK109 cells treated with dTAG-V1, aTAG-2, or binders (GNE-781 + mAP1867-CONHEt) for the indicated duration (1 μM for all compounds). C) Western blot for HA and TFE3 in NONO-TFE3-FKBP^F36V^-HA UOK109 cells treated with aTAG-2 (1 μM) ± competitors (10 μM for mAP1867-CONHEt, 20 μM for GNE-781) after 24 hours. D) RT-qPCR for TFE3 fusion targets (*ANGPTL2, TRIM63*; normalized to *GAPDH*) in a time course following aTAG-2 treatment (50 nM) of NONO-TFE3-FKBP^F36V^-HA UOK109 cells.

**Fig. S5.**
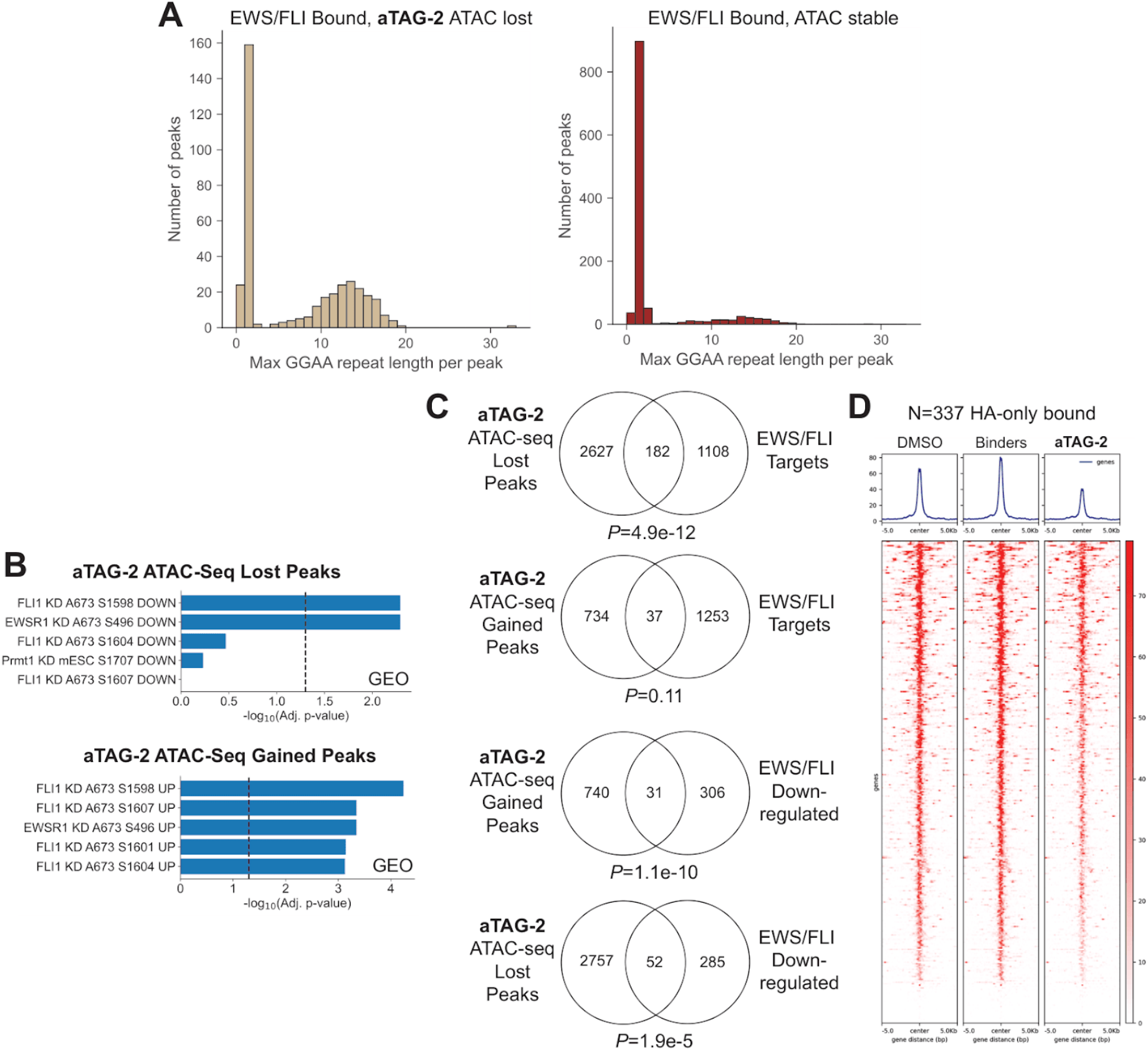
aTAG-2 induces targeted chromatin accessibility changes. FKBP^F36V^-HA-EWS/FLI EWS502 cells treated with DMSO, aTAG-2, or binders (mAP1867-CONHEt + GNE-781) at 5 nM for 16 hours and profiled by ATAC-seq. A) GGAA repeat length in consensus aTAG-2 lost peaks that are EWS/FLI bound (Riggi et al. EWS/FLI bound enhancers^85^) (left) or EWS/FLI-bound ATAC stable peaks (right). B) Enrichr gene ontology analysis on unique genes corresponding to top: consensus aTAG-2 lost ATAC-seq peaks, bottom: consensus aTAG-2 gained ATAC-seq peaks (GEO gene set database). C) Venn diagram of unique genes corresponding to consensus aTAG-2 lost/gained ATAC-seq peaks and EWS/FLI target gene sets from Kinsey et al. (upregulated or downregulated). *P*-values calculated by hypergeometric test. D) ATAC-seq heatmap across HA-bound, p300-unbound peaks are shown (determined using venn diagram in Fig. 3E).

**Fig. S6.**
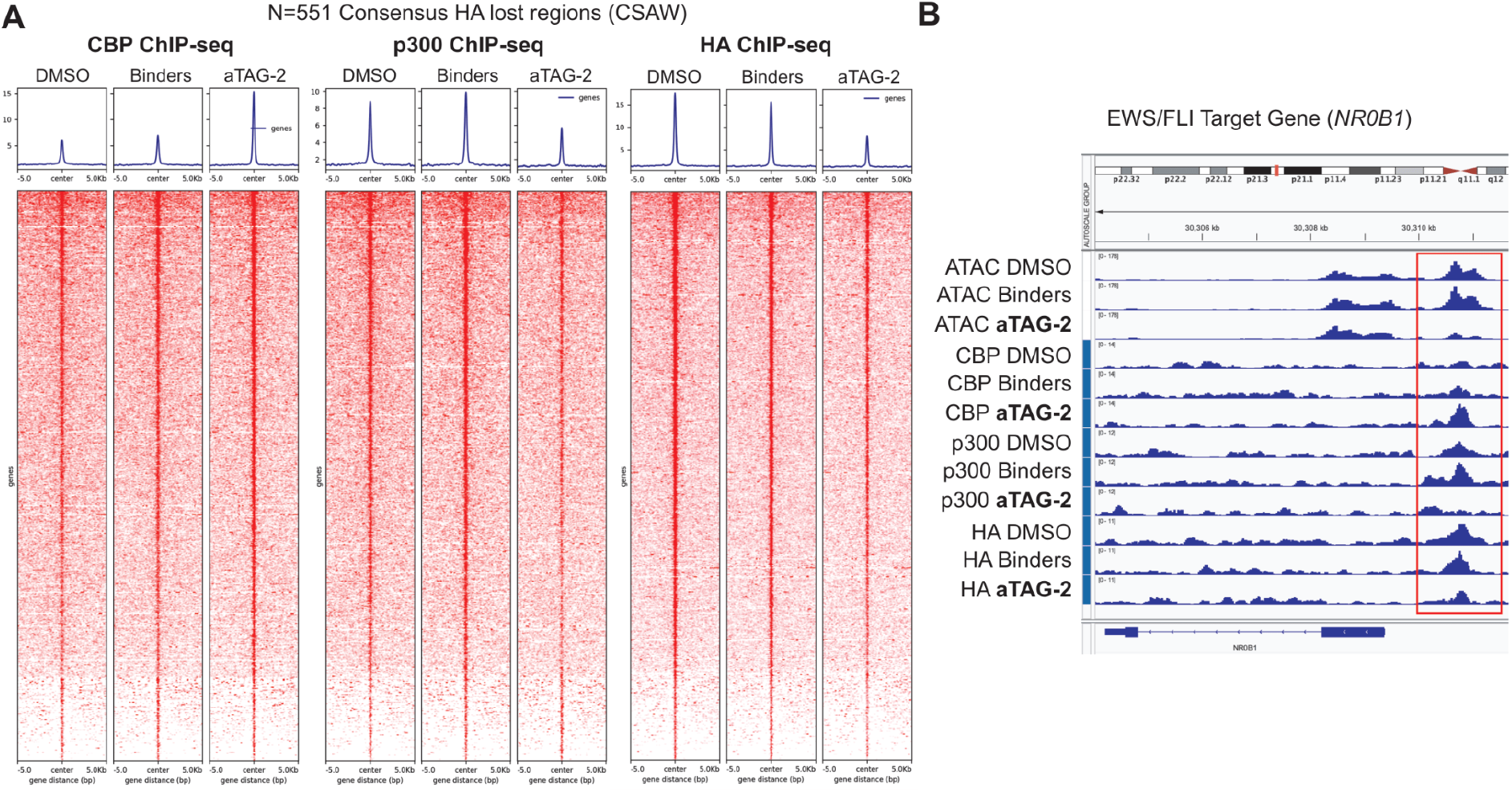
aTAG-2 replaces p300 with CBP at EWS/FLI bound sites. A) FKBP^F36V^-HA-EWS/FLI EWS502 cells treated with DMSO, aTAG-2, or binders (mAP1867-CONHEt + GNE-781) at 10 nM for 16 hours and profiled by CBP, p300, and HA ChIP-seq. P300 and CBP occupancy heatmaps at consensus aTAG-2 HA-EWS/FLI lost regions are shown (N=551, csaw-based differential binding). B) IGV tracks showing changes in ATAC, CBP, p300, and HA occupancy at the best characterized EWS/FLI target gene *NR0B1*.

**Fig. S7.**
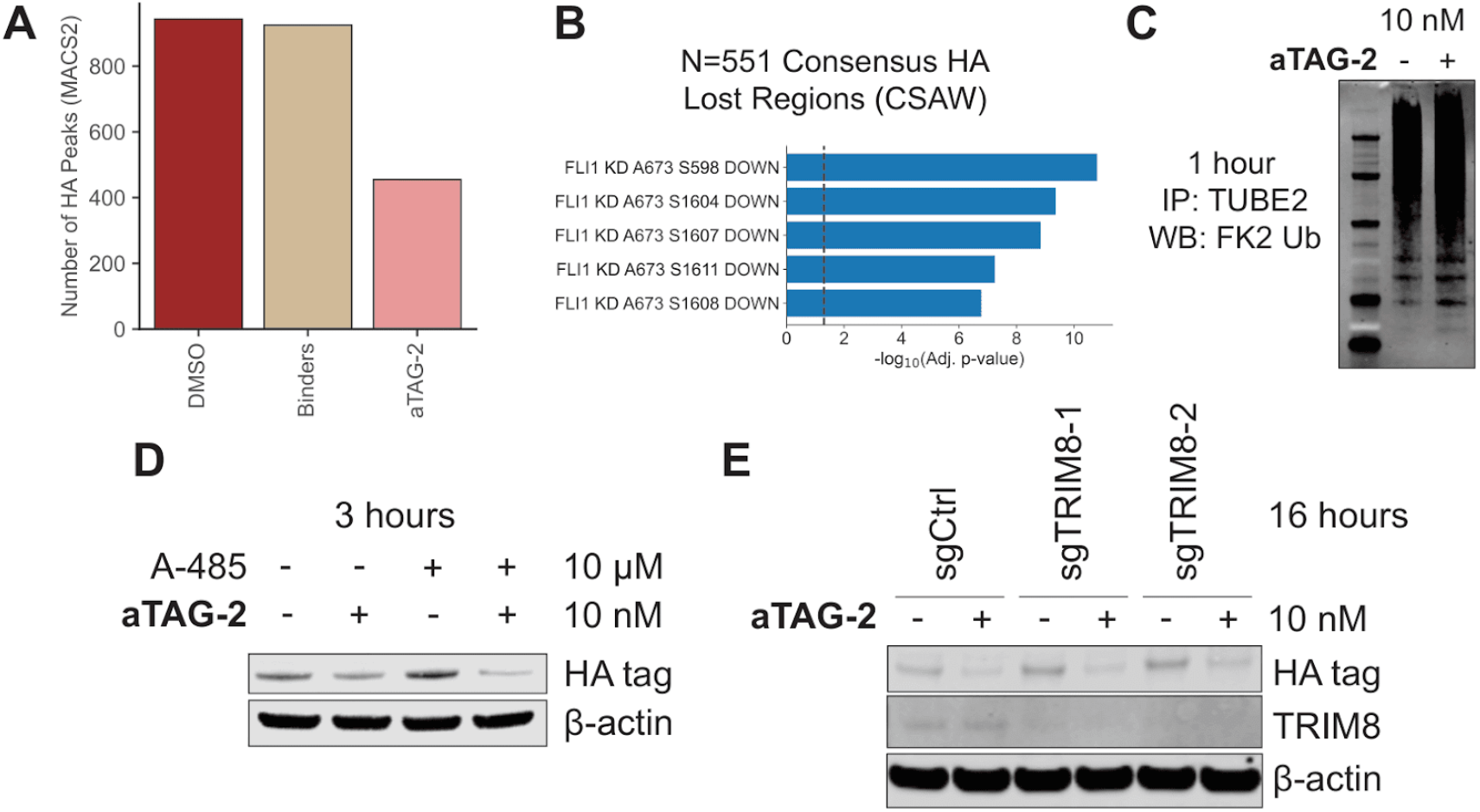
aTAG-2 degrades FKBP^F36V^-HA-tagged EWS/FLI on chromatin. A) FKBP^F36V^-HA-EWS/FLI EWS502 cells treated with DMSO, aTAG-2, or binders (mAP1867-CONHEt + GNE-781) at 10 nM for 16 hours and profiled by HA ChIP-seq. The number of HA peaks in MACS2-based peak calling is plotted between the conditions. B) Gene set enrichment analysis (Enrichr, GEO database) on unique genes corresponding to aTAG-2 HA-EWS/FLI consensus lost regions (N=551, csaw-based differential binding). C) Western blot for total ubiquitin (FK2) in FKBP^F36V^-HA-EWS/FLI EWS502 cells treated with aTAG-2 at 10 nM for 1 hour following TUBE2 (ubiquitin) pulldown, in the presence of MG-132 (10 μM). Loading control for Fig. 4F. D) Western blot for HA-tagged EWS/FLI in FKBP^F36V^-HA-EWS/FLI EWS502 cells treated with aTAG-2 at 10 nM for 3 hours in the presence of p300/CBP histone acetyl transferase inhibitor A-485 (10 μM). E) Western blot for HA-tagged EWS/FLI and TRIM8 in FKBP^F36V^-HA-EWS/FLI EWS502 cells ± dox-inducible TRIM8 sgRNAs treated with aTAG-2 at 10 nM for 16 hours.

